# Fascin-induced bundling protects actin filaments from disassembly by cofilin

**DOI:** 10.1101/2023.05.19.541460

**Authors:** Jahnavi Chikireddy, Léana Lengagne, Rémi Le Borgne, Hugo Wioland, Guillaume Romet-Lemonne, Antoine Jégou

## Abstract

Actin filament turnover plays a central role in shaping actin networks, yet the feedback mechanism between network architecture and filament assembly dynamics remains unclear. The activity of ADF/cofilin, the main protein family responsible for filament disassembly, has been mainly studied at the single filament level. Here, we report that fascin, by crosslinking filaments into bundles, strongly slows down filament disassembly by cofilin. We show that this is mainly due to a slower nucleation of the first cofilin clusters, which occurs up to 100-fold slower on large bundles compared to single filaments. In contrast, severing at cofilin cluster boundaries is unaffected by fascin bundling. After the nucleation of an initial cofilin cluster on a filament of a bundle, we observe the local removal of fascin. Surprisingly, the nucleation of cofilin clusters on adjacent filaments is highly enhanced, locally. We propose that this inter-filament cooperativity in cofilin binding arises from the local propagation of the cofilin-induced change in helicity from one filament to the other filaments of the bundle. Taken together, these observations reveal the molecular events explaining why, despite inter-filament cooperativity, fascin crosslinking protects actin filaments from cofilin-induced disassembly. These findings highlight the important role played by crosslinkers in organizing actin networks and modulating the activity of other regulatory proteins.

## Introduction

Cells assemble a variety of actin filament networks to perform many fundamental cellular functions (Chalut & Paluch, 2016). Importantly, actin filament turnover needs to be tightly controlled for these networks to be functional and to timely adapt to external chemical and mechanical cues (Lappalainen *et al*, 2022; Blanchoin *et al*, 2014). Typically, actin filaments that polymerize in lamellipodia or the actin cortex are renewed within a few seconds (Fritzsche *et al*, 2013; Lai *et al*, 2008), while filaments composing stress fibers are much more stable, turning over several minutes (Campbell & Knight, 2007; Saito *et al*, 2022; Valencia *et al*, 2021).

These actin networks are exposed to disassembly factors, among which proteins of the ADF/cofilin family are the main players (Svitkina & Borisy, 1999; Hotulainen *et al*, 2005). ADF/cofilin (hereafter cofilin) activity has been extensively characterized at the single actin filament level. It is known to induce actin filament fragmentation (Carlier *et al*, 1997; Blanchoin & Pollard, 1999), as well as filament depolymerization from both ends (McGough *et al*, 1997; McCullough *et al*, 2008; Suarez *et al*, 2011; Wioland *et al*, 2017; Schramm *et al*, 2017). Cofilin-induced severing is a complex mechanism that involves several reaction steps. Cofilin binds preferentially to ADP-actin filament segments rather than to ‘younger’ ADP-Pi-rich segments (Maciver *et al*, 1991; Carlier *et al*, 1997; Suarez *et al*, 2011). The binding of cofilin is cooperative, as cofilin molecules bind with a higher affinity to a site adjacent to a cofilin-occupied site along a filament, creating cofilin clusters (De La Cruz, 2005). Moreover, cofilin binding locally shortens the helical pitch of filaments (Galkin *et al*, 2001). Filament severing occurs at cluster boundaries, and 4 times more frequently at the boundary on the pointed end side of the cluster (Suarez *et al*, 2011; Gressin *et al*, 2015; Wioland *et al*, 2017). Recent CryoEM observations have revealed, with unprecedented details, that the change of filament and actin subunits conformations upon cofilin binding does not propagate more than one actin subunit away from the cofilin boundary (Huehn *et al*, 2020). Reaction rates associated with cofilin cluster nucleation, growth and severing have been measured for the three mammalian ADF/cofilin isoforms on α-skeletal and cytoplasmic β- and γ-actin filaments (Wioland *et al*, 2017, 2019a). Importantly, the ability of cofilin to apply a torque on twist-constrained filaments increases the severing rate by up to 2 orders of magnitude (Wioland *et al*, 2019b). This last result strongly suggests that the geometrical constraints imposed by crosslinkers in filament networks should enhance the ability of cofilin to disassemble these networks.

In cells, actin filaments can be crosslinked by a large array of crosslinker proteins that vary in size, affinity for the side of filaments, and in the way they decorate filaments (Blanchoin *et al*, 2014; Gallop, 2019; Rajan *et al*, 2023). As a consequence, filament crosslinkers are major contributors of actin network architectures, by specifically tuning their geometrical and mechanical properties (Claessens *et al*, 2006; Ma & Berro, 2018; Freedman *et al*, 2019; Banerjee *et al*, 2020; Lieleg *et al*, 2010). One special case of filament crosslinking is parallel filament bundling. Such actin filament organization is observed in filopodia and microspikes that emerge from the front of lamellipodia (Vignjevic *et al*, 2006), as well as in microvilli and stereocilia.

Previous studies have revealed that fascin is the main filament crosslinker in filopodia and microspikes (Adams & Schwartz, 2000; Vignjevic *et al*, 2006; Faix *et al*, 2009; Jacquemet *et al*, 2019; Damiano-Guercio *et al*, 2020). Fascin is a monomeric protein that arranges parallel filaments into bundles, with an hexagonal packing (Aramaki *et al*, 2016; Shin *et al*, 2009). In between two filaments of these bundles, fascin binds regularly, every half-pitch (36 nm) along actin filaments (Edwards *et al*, 1995; Ishikawa *et al*, 2003; Jansen *et al*, 2011; Aramaki *et al*, 2016), undertwists filaments by 1° (Shin *et al*, 2009), and quickly turns over (Aratyn *et al*, 2007; Winkelman *et al*, 2016; Suzuki *et al*, 2020). Bundles formed by fascin typically reach a maximum size of typically 15 filaments both in vivo and in vitro (Jansen *et al*, 2011; Breitsprecher *et al*, 2011; Aramaki *et al*, 2016; Atherton *et al*, 2022; Hylton *et al*, 2022). This limit in size is thought to arise from the constraint of aligning the fascin binding sites among the actin filaments (Claessens *et al*, 2008).

So far, to our knowledge, only one study has addressed in vitro the impact of fascin bundling on cofilin disassembly activity (Breitsprecher *et al*, 2011). Using both bulk pyrene and TIRF microscopy assays, Breitsprecher and colleagues proposed that fascin, by preventing the relaxation of the cofilin-induced torque, favored filament severing by cofilin. However, the absence of direct visualization of cofilin binding and severing events on filament bundles prompted us to further investigate the molecular details of the disassembly of fascin-induced filament bundles by cofilin.

Here, using purified proteins, we show that cofilin binds and fragments filament bundles slower than single actin filaments (Fig. 1 & 2). We report that cofilin cluster nucleation and growth rates on fascin-induced filament bundles are reduced compared to what is observed on single filaments (Fig. 3 & 4). We further show that fascin is removed as cofilin clusters expand (Fig. 5). Strikingly, we reveal and quantify an inter-filament cooperativity mechanism where, after the creation of a first cofilin cluster, the subsequent nucleation of cofilin clusters on adjacent filaments within a bundle is strongly enhanced (Fig. 6). We propose a model where filament crosslinking allows the cofilin-induced local change of helicity of one filament to be transmitted to other filaments (Fig. 6). Last, numerical simulations integrating all the cofilin reaction rates determined experimentally recapitulate the observed fragmentation of both twist-unconstrained and twist-constrained actin filament bundles (Fig. 7). Overall, our in vitro results provide novel molecular insights into how cofilin-induced actin network disassembly is affected by filament crosslinking and organization in cells.

**Figure 1.**
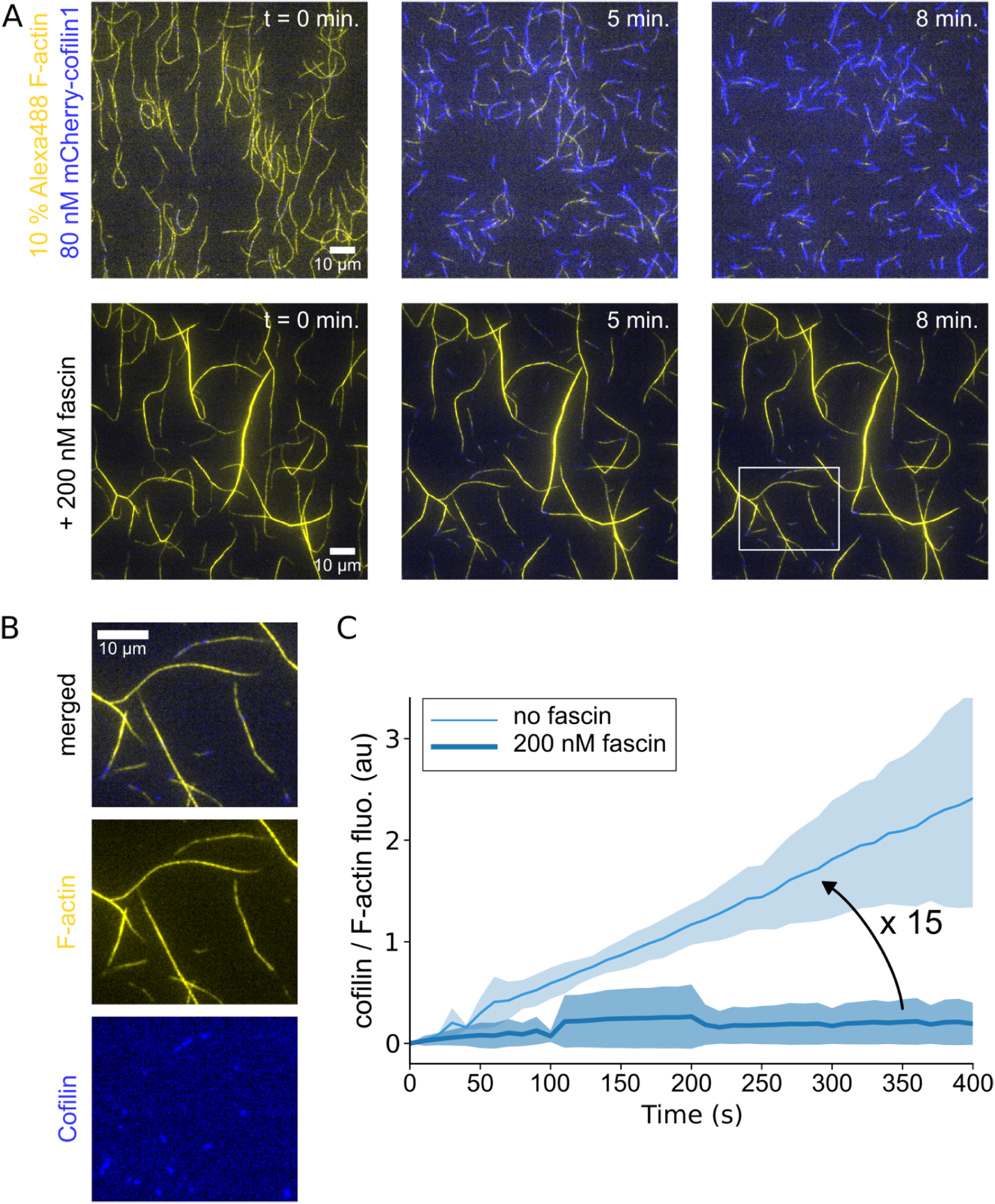
Fascin-induced actin filament bundling slows down the recruitment of cofilin. **A.** 10% Alexa-488 labeled actin filaments, grown and aged for 15 minutes from surface anchored seeds, in the absence or presence of 200 nM human-fascin1, in a buffer containing 0.2% methylcellulose, are subsequently exposed to 80 nM mCherry-cofilin1 at time 0 (see Methods). **B.** Fluorescence image of the rectangular area shown in A, in the presence of fascin, with the different channels shown separately. **C.** Fluorescence intensity of bound cofilin, normalized by the amount of F-actin, as a function of time. Both curves increase roughly linearly. Linear fitting indicates that cofilin binding to single actin filaments is 15 times faster than on fascin-induced actin filament bundles (N=3 independent experiments). Shaded areas represent standard deviations.

**Figure 2.**
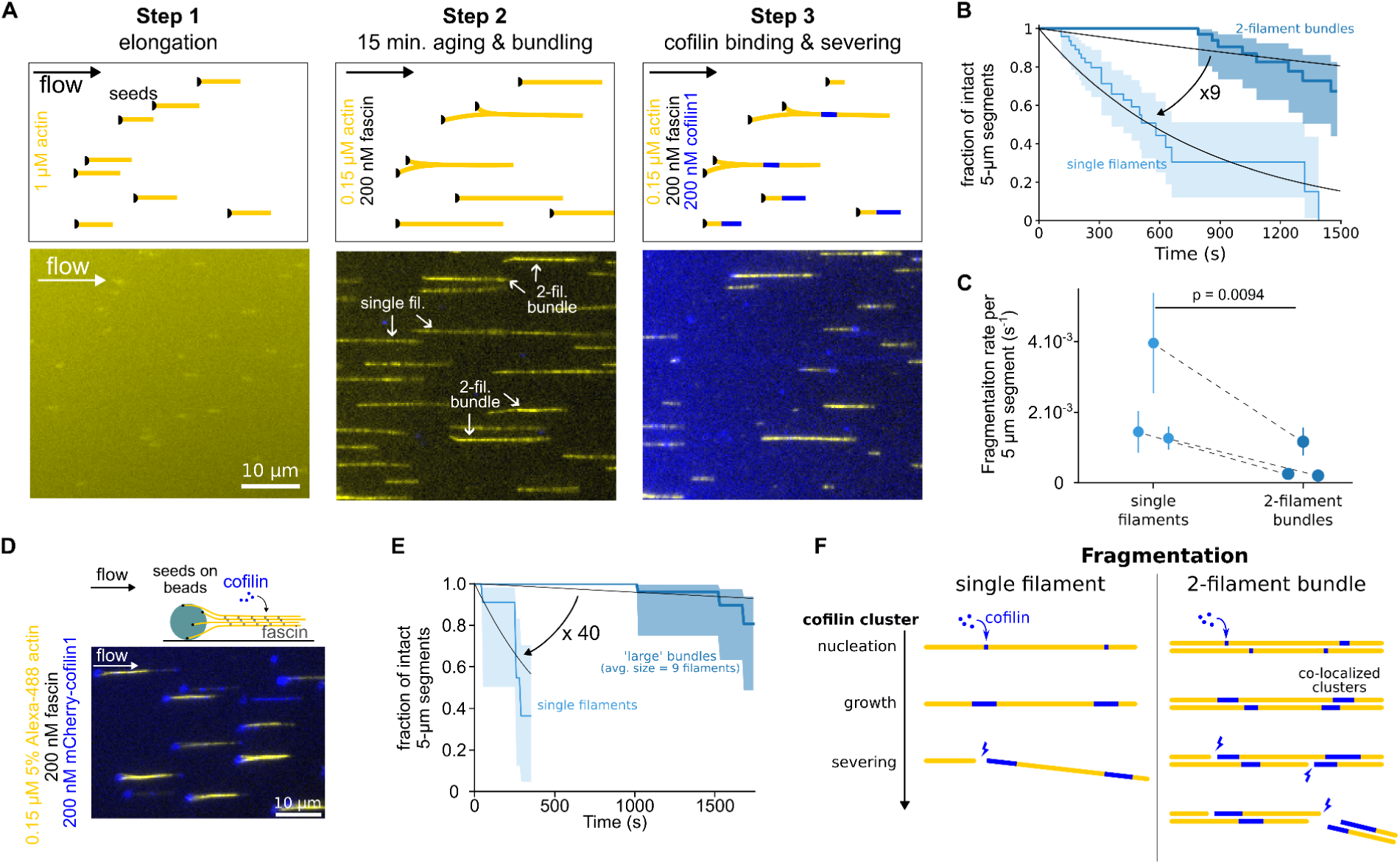
Cofilin fragments fascin-induced bundles slower than single filaments. **A.** Schematics of the sequential steps to investigate fascin-induced 2-filament bundle fragmentation by cofilin. Inside a microfluidic chamber, actin filaments were grown from randomly positioned surface-anchored seeds and aligned by the flow (step 1). Once elongated, filaments were aged and allowed to form 2-filament bundles in the presence of fascin and actin for 15 minutes (step 2). They are then exposed to cofilin, actin and fascin (step 3). Note that isolated filaments did not bundle and were used as reference single filaments for side-by-side comparison. **B.** Result from a typical experiment showing the fragmentation, over time, of single actin filaments (n = 53) and 2-filament bundles (n = 47) upon exposure to 200 nM cofilin, 0.15 µM actin and 200 nM fascin. 95% confidence intervals are shown as shaded surfaces. There is a ∼ 9-fold difference in the rate of decay of those two populations, as obtained by single exponential fits (lines). **C.** Fragmentation rates of single actin filaments and 2-filament bundles when exposed to 200 nM cofilin. Rates are obtained from exponential fits as shown in panel B. Dashed lines indicate paired data points from single filament and bundle populations acquired simultaneously in the same microfluidics chamber (N = 3 independent experiments; n= 53, 21, 29 single filaments; n = 47, 21, 30 2-filaments bundles). Rates and error bars are obtained from exponential fits as shown in panel B. **D.** Actin filaments were polymerized from spectrin-actin seeds adsorbed to micron-sized glass beads to create larger bundles, in an experiment otherwise similar to the one shown in panel A. Note that non-productive spectrin-actin seeds on beads are targeted by cofilin (blue), as revealed by the fluorescence on the surface of the bead. **E.** Survival fractions of intact 5-µm long segments of single actin filaments (n = 12) or filament bundles (n = 30, average size 9.3 (± 3.2) filaments per bundle), upon exposure to 200 nM cofilin and 200 nM fascin, as a function of time. 95% confidence intervals are shown as shaded surfaces. There is a ∼ 40-fold difference in the rates at which these two populations decrease, as obtained by single exponential fits (lines). **F.** Schematics of the reactions that lead to cofilin-induced severing. The fragmentation of single filaments (left) results from the severing of one cofilin cluster, whereas the fragmentation of 2-filament bundles (right) requires the severing of two ‘co-localized’ cofilin clusters, one on each filament.

**Figure 3.**
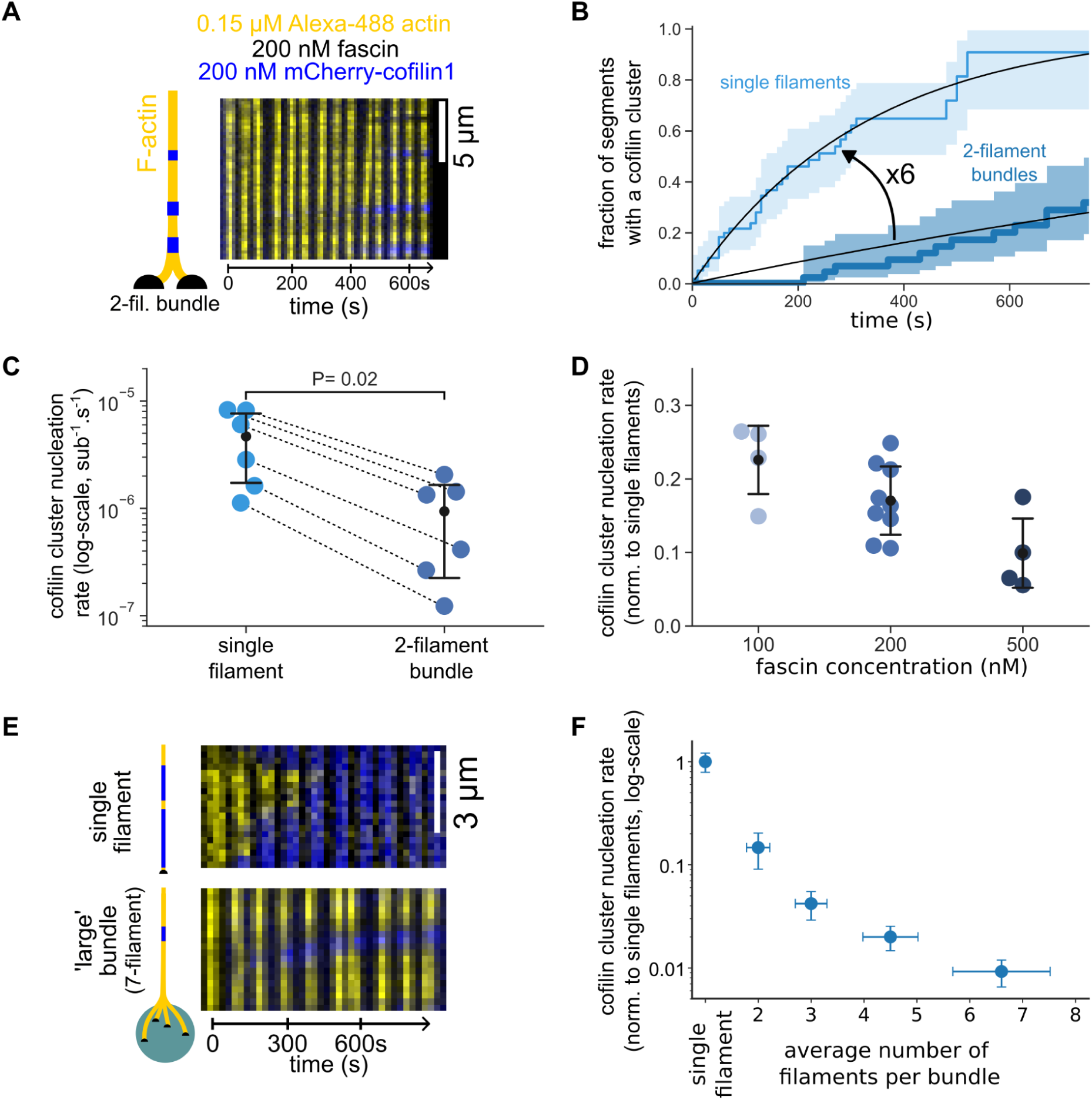
Cofilin cluster nucleation is slowed down by fascin-induced filament bundling. **A.** Time-lapse images of a 2-filament bundle (yellow) with nucleation of cofilin clusters (blue). **B.** The fraction of 5-µm segments of single filaments (light blue, n= 63 segments) or 2.5-µm segments of 2-filament bundles (dark blue, n= 47 segments) where at least one cofilin cluster has nucleated, over time. 95% confidence intervals are shown as shaded surfaces. Black lines are single exponential fits. **C.** Cofilin cluster nucleation rates per cofilin binding site along actin filaments (log-scale), measured from 6 independent experiments (n > 30 segments for each population), in the presence of 200 nM cofilin, 200 nM fascin and 0.15 µM actin. For each condition, the black dot represents the average and the error bars represent the standard deviation of the distribution. Rates were obtained from exponential fits as shown in panel B. Dashed lines indicate paired data points from populations acquired simultaneously in the same microchamber. The paired data shows consistently a 6-fold difference in nucleation rates. The p-value is from a paired t-test. **D.** Impact of fascin concentration on the cofilin cluster nucleation rate on 2-filament bundles, normalized by the rate on single filaments (n = 4, 9 and 4 independent experiments at 100, 200 and 500 nM fascin respectively, with >20 segments analyzed for each experiment). Rates were derived as shown in panel B. All conditions with 200 nM cofilin and 0.15 µM actin. For each fascin concentration, the error bar represents the standard deviation of the distribution. **E.** Timelapse images showing the appearance of cofilin clusters (blue) on either a single actin filament or a 7-filament bundle (yellow) grown from a micrometer-size glass bead, in conditions similar to panel A. **F.** Impact of bundle size on the cofilin cluster nucleation rate (per cofilin binding site), normalized by the rate on single filaments, when exposed to 200 nM cofilin, 200 nM fascin and 0.15 µM actin (N = 1 experiment, with 22, 10, 20, 40 and 25 segments analyzed for single filaments, and bundles of size 2, 3, 4-5, and 5-8 filaments, respectively). Error bars are standard deviations for bundle size and the normalized cofilin nucleation rates.

**Figure 4.**
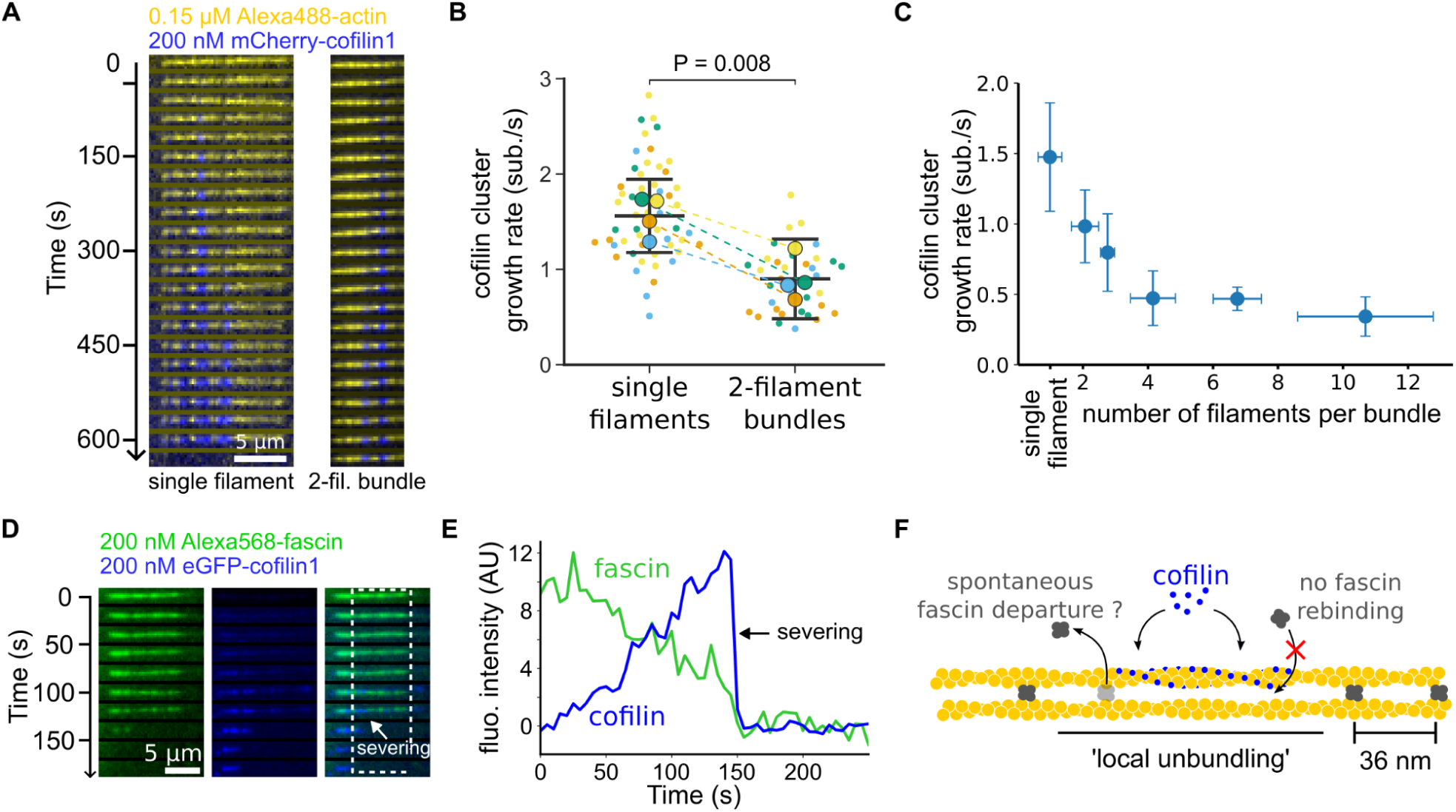
Cofilin and fascin compete to bind filaments in bundles. **A.** Time-lapse images showing the growth of a cofilin cluster (blue) on a 2-filament bundle (yellow). **B.** Cofilin cluster growth rates on single filaments and 2-filament bundles, in the presence of 200 nM cofilin, 200 nM fascin and 0.15 µM Alexa488(10%)-G-actin (N = 4 repeats, each of a different color, with at least 10 cofilin clusters observed in each condition). Small data points are individual measurements (one per cluster) and the large points are the averages per repeat. Dashed lines indicate paired averages from populations acquired simultaneously in the same microchamber. The p-value is from a paired t-test. **C.** Cofilin cluster growth rates as a function of bundle size (n=11, 4, 6, 7, 6 and 9 cofilin clusters analyzed on single filaments and bundles of an average size of 2, 3, 4, 7 and 11 filaments, respectively). Error bars are standard deviations. **D.** Time-lapse images showing a 2-filament bundle with Alexa568-fascin (green) and exposed to eGFP-cofilin1 (blue). **E.** Fluorescence intensity of fascin and cofilin, integrated over the region shown in panel D (dashed rectangle), as a function of time. **F.** Schematics of the irreversible departure of fascin, caused by a cofilin cluster growing on a 2-filament bundle.

**Figure 5.**
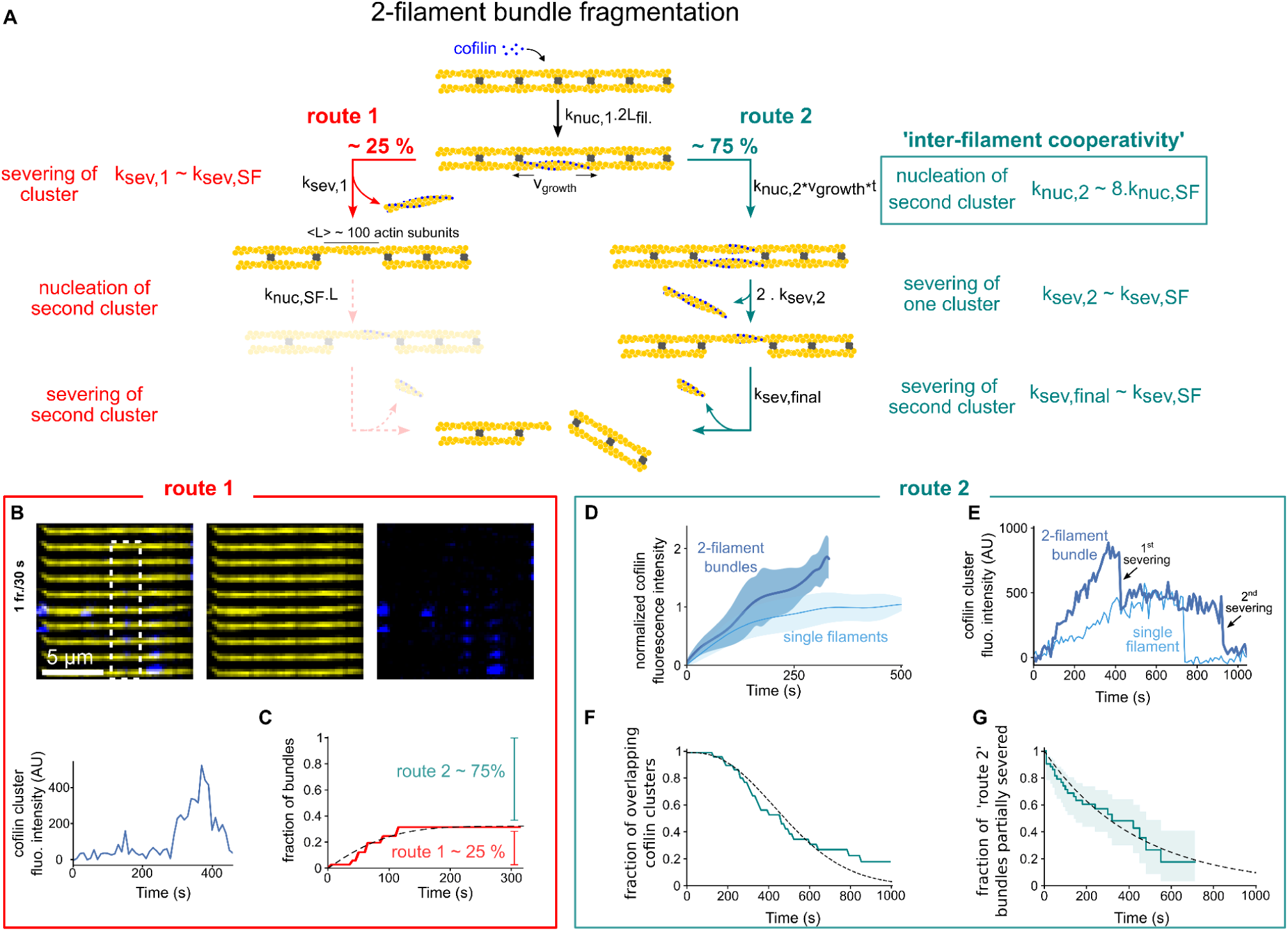
Inter-filament cooperativity leads to efficient bundle fragmentation. **A.** schematic representation of the two routes leading to the fragmentation of a 2-filament actin bundle, after the first cofilin cluster has nucleated, at a rate k_nuc,1_.2L_fil_, where L_fil_ is the segment length of each filament of the bundle. For route 1, the initial cofilin cluster severs at its two boundaries (see main text), at a rate k_sev,1_, before another cofilin cluster is nucleated on the other filament of the bundle, in the region facing the first cofilin cluster. Subsequently, the nucleation of a cluster on the remaining filament, followed by its severing, leads to bundle fragmentation, but these two final steps were never observed experimentally (shaded steps). For route 2, a cofilin cluster nucleates on the second filament in the region facing the first cofilin cluster before the latter severs, leading to the presence of two clusters in the same region. The sequential severing of the two clusters fragments the bundle. **B.** An example illustrating the route 1 type of events: a single cofilin cluster (blue) severs a region of one filament in a 2-filament bundle, while no other cofilin cluster has nucleated on the adjacent filament. The graph shows the intensity of the cofilin cluster in the dashed rectangle, as a function of time. **C.** Fraction of bundles for which the first cofilin cluster severs, leaving behind no detectable cofilin, as a function of time. The fit of the experimental curve (see Methods) yields k_sev,1_ = 3.10^-3^ s^-1^ and k_nuc,2_ = 1.10^-4^ sub^-1^.s^-1^ (n = 71 events). The dashed line shows the best fit, using a chi-square minimization procedure (see Methods). Three additional experiment repeats are shown in Supp. Fig. 10. In total, N = 4 independent experiments, with n = 101, 71, 17, 28 events, yield the following average values of k_sev,1_=1.5.10^-3^ s^-1^ and k_nuc,2_ = 4.7.10^-5^ sub^-1^.s^-1^. **D.** Average fluorescence intensity of cofilin clusters over a 1 µm wide segment on single filaments (light blue, n = 10) and on 2-filament bundles (dark blue, n = 29), as a function of time, in the presence of 200 nM fascin and 200 nM cofilin, normalized by the maximum fluorescence intensity on single filaments. **E.** Examples of cofilin intensity traces on a single filament (light blue) and a 2-filament bundle (dark blue) as a function of time. The drop in intensity for the single filament is accompanied by the fragmentation of the single filament. For the 2-filament bundle, a drop in the intensity to a lower value reveals the existence of 2 ‘co-localized’ cofilin clusters. The second drop in intensity is accompanied by the complete fragmentation of the bundle. **F.** Fraction of co-localized cluster regions that have not yet had a severing event, versus time (n = 31 events). The dashed line shows the best fit by a computed curve (see Methods), using a chi-square minimization procedure. 3 experimental repeats, with n = 31, 33, 39 events each, yield an average k_sev,2_ = 1.06 (± 0.2).10^-3^ s^-1^ (± standard deviation). **G.** Fraction of cofilin clusters remaining after the first severing event that have not yet had the second severing event, versus time (n = 53 events, pooled from 4 independent experiments). Fit of the experimental curve by a single exponential yields k_sev,final_ = 2.25 (± 0.6).10^-3^ s^-1^ (± 95% confidence interval).

**Figure 6.**
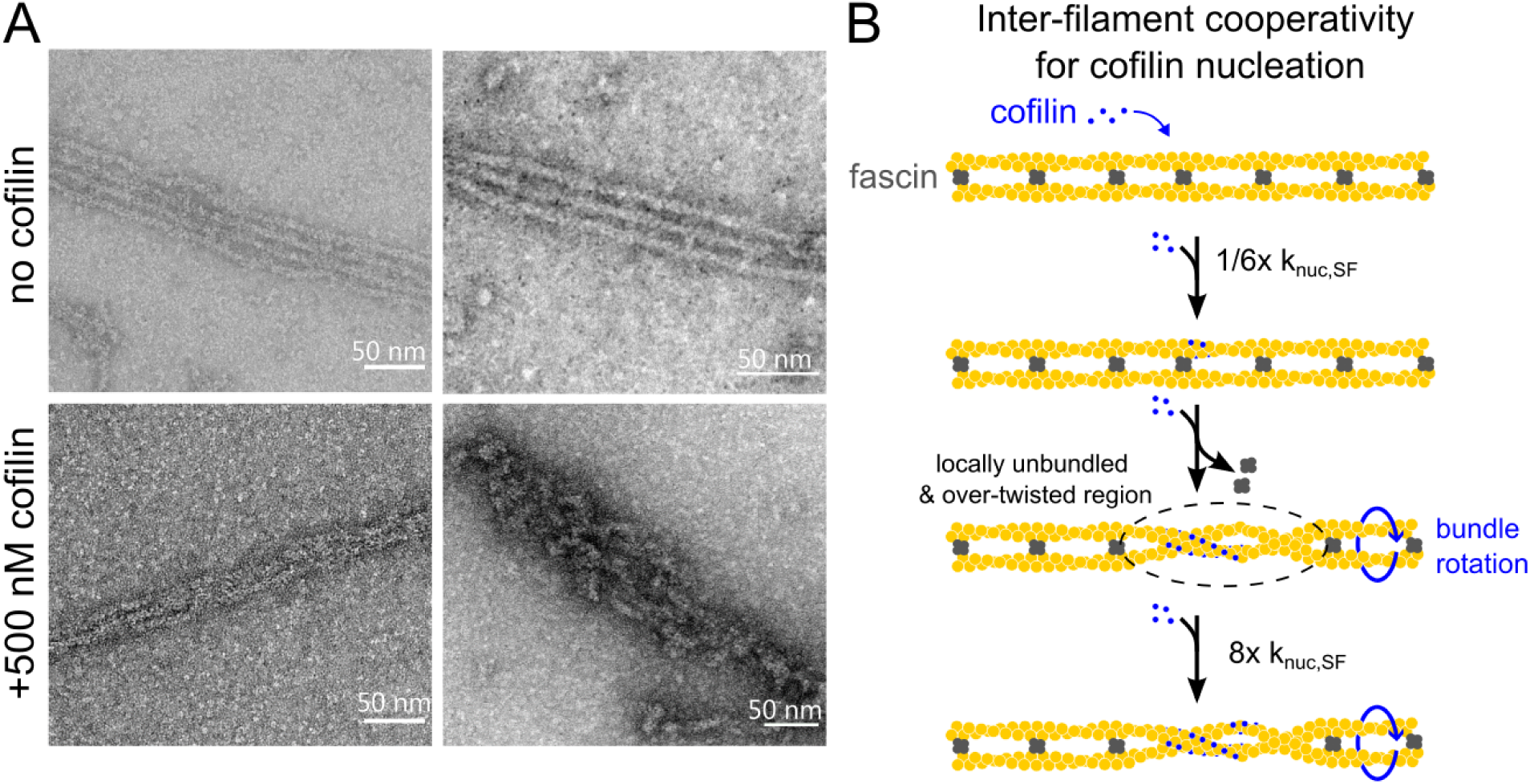
Cofilin locally triggers inter-filament twisting of fascin-induced bundles. **A.** Negative-staining electron micrographs of fascin-induced bundles in the presence or absence of 500 nM cofilin, incubated for 3 minutes before the solution was adsorbed on a freshly glow-discharged carbon-coated copper grid. **B.** Schematics of the inter-filament cooperative twisting model. For a fascin-induced 2-filament bundle, a first cofilin cluster is nucleated on one of the filaments and starts growing, preventing fascin from binding locally. Local over-twisting caused by cofilin decoration is transmitted to the adjacent filament in the region devoid of fascin. This favors the binding of cofilin on the undecorated filament: the nucleation rate of the second cofilin cluster is 48-fold higher than for the initial cofilin cluster.

**Figure 7.**
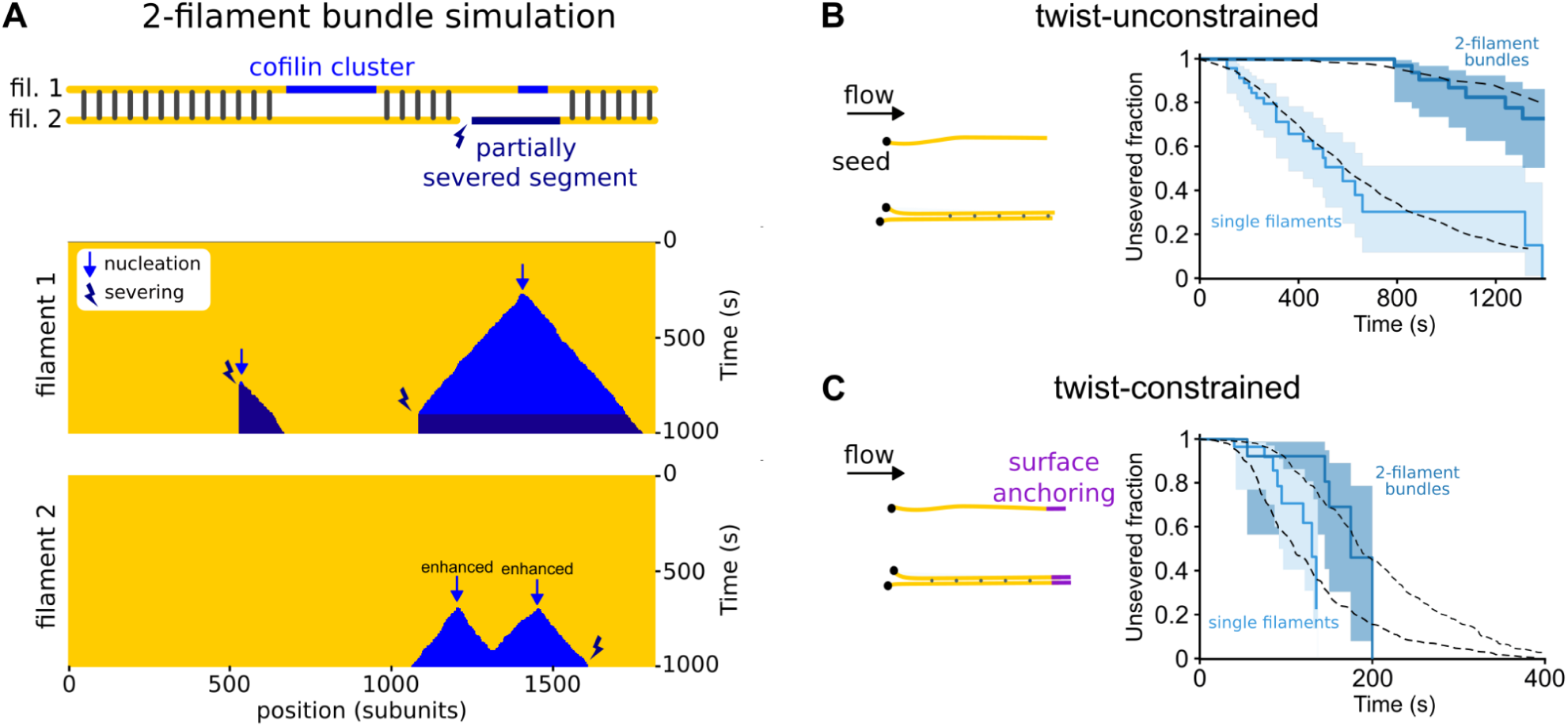
Constraining 2-filament bundles in twist highly enhances their fragmentation. **A.** (top) Schematics illustrating the numerical simulations of two filaments of 5-µm segments (yellow) interconnected by fascin (dark gray), where cofilin clusters (light blue) can nucleate and sever filaments (dark blue). (bottom) Kymographs of two interconnected simulated filaments, showing cofilin cluster nucleation (arrow) and severing (thunderbolt) events on each filament. The kymographs stop when the severing events on the two filaments result in the fragmentation of the bundle. Numerical values of reaction rates are summarized in Supp. Table S1. **B,C.** Fraction of unsevered 5-µm segments that are (B) twist-unconstrained or (C) twist-constrained by being doubly attached to the glass surface, for single actin filaments (light blue, n = 53, 34 for filaments unconstrained and constrained in twist resp.) or 2-filament bundles (dark blue, n = 47, 16 for bundles unconstrained and constrained in twist resp.) upon exposure to 200 nM cofilin and 200 nM fascin, as a function of time. 95% confidence intervals are shown as shaded surfaces. Dashed lines correspond to the results obtained from numerically simulated segments (n = 200 for twist-unconstrained, 50 for twist-constrained segments), using experimentally determined rates and considering no inter-filament cooperativity in twist-constrained bundles (see text).

## Results

All experiments were performed at 25°C, in a buffer at pH 7.4, with 50 mM KCl, 1 mM MgCl_2_, using rabbit α-skeletal actin, recombinantly expressed mouse cofilin-1 (cofilin from here on) and human fascin-1 (fascin from here on) (see Methods).

### Fascin-induced bundles are protected from cofilin binding

In order to investigate the overall activity of cofilin on fascin-induced filament bundles, we first performed experiments in so-called ‘open chambers’, in a buffer supplemented with 0.2% methylcellulose (see Methods). Actin filaments were polymerized from surface-anchored spectrin-actin seeds, and exposed to 200 nM fascin as they grew. This fascin concentration allowed adjacent filaments to rapidly form stable bundles (Supp. movie 1), reaching on average 10 (+/- 5) filaments per bundle, as estimated per their actin fluorescence intensity. Actin filaments were then exposed to actin at critical concentration (∼ 0.15 µM) and 200 nM fascin for 15 minutes, so that filaments remained bundled, and maintained a constant length as they became ADP-actin filaments. Upon addition of 80 nM mCherry-cofilin1, cofilin clusters appeared only very sparsely along filament bundles and the network did not disassemble over the course of 10 minutes (Fig. 1A). In contrast, in control experiments without fascin, single actin filaments were readily targeted by cofilin, leading to their rapid fragmentation and disassembly (Supp. movie 2).

By quantifying the increase of cofilin fluorescence, normalized by the F-actin fluorescence, we observed that the binding rate of cofilin was ∼ 15-fold slower on fascin-induced bundles than on single actin filaments (Fig. 1B,C). Interestingly, larger bundles recruit less cofilin than smaller ones (∼ 2.5 times less cofilin bound per F-actin for bundles of 10 (± 4) filaments than of 3 (± 0.8) filaments, after 8 minutes, Supp. Fig. 1).

### Cofilin fragments 2-filament bundles more slowly than single filaments

In order to investigate the molecular mechanisms of actin filament bundle disassembly by cofilin in more controlled conditions, we performed experiments using microfluidics (Jégou *et al*, 2011; Wioland *et al*, 2022). We first sought to quantify cofilin activity on 2-filament bundles, the smallest bundle unit that can be assembled by fascin (Fig. 2A, Supp. movie 3). Briefly, actin filaments were elongated from randomly positioned spectrin-actin seeds anchored on the glass surface of a microfluidic chamber, to reach typically 10 µm in length. Upon exposure to 200 nM fascin, filaments bundled together, forming a majority of 2-filament bundles, as revealed by their actin fluorescence intensity. Larger bundles were discarded from the analysis for this type of assay. Filaments that were far from adjacent filaments could not form bundles, thereby providing a reference population of single filaments for the analysis. Filaments were aged for 15 minutes to become ADP-actin filaments in the presence of fascin and actin. We verified that fascin crosslinking did not appear to significantly slow down phosphate release in actin filaments (Supp. Fig. 2), which otherwise would have affected cofilin binding on bundles compared to single filaments. Both bundles and single filaments were then exposed to 200 nM mCherry-cofilin-1, in the presence of fascin and actin to maintain filament bundling and filament length. We quantified the fraction of intact 5-µm long segments for single actin filaments and for 2-filament bundles, as a function of time upon exposure to cofilin. We observed a ∼ 9-fold slower fragmentation for 2-filament bundles (Fig. 2B, C).

To specifically investigate cofilin-induced fragmentation on larger bundles, we grew several actin filaments from spectrin-actin seeds attached to micrometer-size beads (in average, 10 filaments per bundle, Supp. Fig. 4A) (Fig. 2D,E). Cofilin fragmentation on those large bundles was strongly reduced, by 40-fold compared to single actin filaments present in the same chamber (Fig. 2F).

Fragmentation of a 2-filament bundle requires the severing of two co-localized cofilin clusters, one on each filament, facing each other (Fig 2F). On its own, this constraint contributes to making the fragmentation of a filament bundle slower than that of a single filament. However, this does not explain why cofilin decoration appears to be slower on bundles than on single filaments (Fig 1). Further, one may expect the severing rate per cofilin cluster to be faster in bundles than single filaments, because of potential constraints on the twist imposed by filament bundling (Wioland *et al*, 2019b; Breitsprecher *et al*, 2011). To understand the mechanism responsible for the slower fragmentation of bundles, we thus decided to quantify the impact of bundling on the different molecular reactions that lead to fragmentation.

### Cofilin clusters nucleate slower on actin filaments bundled by fascin

To better understand how cofilin disassembles filament bundles, we next sought to quantify cofilin cluster nucleation, which is the first step in cofilin-induced filament disassembly. First, we measured the impact of fascin on the nucleation of cofilin clusters on single filaments. While fluorescently-labeled fascin is easily detected on bundles formed from nearby filaments (Aratyn *et al*, 2007; Winkelman *et al*, 2016; Suzuki *et al*, 2020), we could not detect the presence of fascin on single actin filaments, even at micromolar concentrations, as previously reported (Winkelman *et al*, 2016; Suzuki *et al*, 2020). Nonetheless, exposing single actin filaments to increasing fascin concentrations gradually decreased the nucleation rate of cofilin clusters, by up to 2-fold in the presence of 1 µM fascin, compared to the absence of fascin (Supp. Fig. 3). At 200 nM fascin, our reference fascin concentration in this study, cofilin binding to filaments was not measurably affected.

In spite of having two filaments, thus twice as many potential binding sites, the nucleation rate of cofilin clusters per binding site on 2-filament bundles was reduced 6-fold, compared to single actin filaments (Fig. 3A-C). This observed strong reduction in the nucleation rate thus appears to play a key role in protecting bundles from cofilin-induced fragmentation.

We observed that cofilin clusters were homogeneously distributed along the bundle, excluding any effect of the moderate pulling force gradient applied on filament bundles by the microfluidics flow (force range 0-0.5 pN, Supp Fig 4). Higher fascin concentrations further decreased the cofilin cluster nucleation rate on 2-filament bundles compared to single actin filaments, up to 10-fold at 500 nM fascin (Fig. 3D). At saturating fascin concentration, fascin binds every 13 actin subunits (*i.e.* every half-pitch of the actin filament) between 2 actin filaments (Jansen *et al*, 2011; Aramaki *et al*, 2016), leaving potentially 12 cofilin binding sites available between two bound fascins, on each filament. Thus, the observed 6- to 10-fold reduction in the nucleation rate of cofilin clusters cannot be explained by a simple competition between cofilin and fascin to bind on actin filaments. Rather, the slower nucleation may originate from the filament helicity imposed by fascin bundling (Shin *et al*, 2009) (Claessens *et al*, 2008).

We observed that the cofilin nucleation rate, per cofilin binding site, strongly decreased with increasing bundle size (Fig. 3F). When measuring the effect per bundle, the nucleation rate is reduced more than 10-fold per bundle for bundles larger than 5 filaments compared to single filaments (Supp. Fig. 5). This is striking as large bundles harbors more cofilin binding sites than smaller bundles. Overall, these results consistently show that fascin-induced bundling strongly decreases the rate of cofilin cluster nucleation.

### Cofilin cluster growth is slowed down by fascin-induced bundling

How fast cofilin clusters grow and decorate actin filaments is an important aspect of cofilin-induced disassembly. We thus next sought to measure the cofilin cluster growth rate on fascin-induced filament bundles. We observed that cofilin clusters grew at a 1.5-fold lower rate on 2-filament bundles than on single filaments, in the presence of 200 nM (Fig. 4 A,B). This observation is consistent with the slower cofilin cluster nucleation reported above, although the effect appears to be much milder for cluster growth. Surprisingly, increasing fascin concentration did not appreciably decrease the cofilin cluster growth rate further (Supp. Fig 6). Cofilin cluster growth rate decreased with bundle size and seemed to plateau for large bundles (Fig. 4C). Considering that actin filaments are arranged hexagonally in fascin-induced bundles, each filament can potentially be crosslinked to up to 6 filament neighbors, which would correspond to a maximum of 6 fascin proteins bound every 13 actin subunits. Taken together, these observations seem consistent with cofilin binding being slowed, during the growth of a cofilin cluster in a bundle, by the density of fascin bound along the filaments. This density depends mainly on the number of adjacent filaments, and only marginally on the concentration of fascin in solution, in the range we used.

Cofilin binds actin subunits stoichiometrically (Galkin *et al*, 2001). We thus asked whether cofilin-saturated regions could also contain fascin. We observed that fluorescently-labeled fascin was gradually excluded from expanding cofilin clusters on 2-filament bundles (Fig. 4D,E). One possible explanation could be that, as fascin quickly turns over in filament bundles (Aratyn *et al*, 2007; Suzuki *et al*, 2020), the departure of fascin would free space for cofilin and allow clusters to grow. This process seems irreversible as increasing the concentration of fascin in solution did not slow down cofilin cluster growth, as would be expected if both proteins were directly competing for the same or overlapping binding sites (Supp. Fig. 6). Importantly, fascin exclusion from a cofilin-saturated region induces filaments of the bundle to be no longer crosslinked locally (‘local unbundling’, Fig. 4F).

### Inter-filament cooperative nucleation of cofilin clusters

As fascin is locally excluded by the presence of a cofilin cluster on one of the two filaments of the bundle, we next sought to investigate whether this would favor the local nucleation of a second cofilin cluster on the other filament. Plotting the local increase in fluorescence intensity of cofilin along 2-filament bundles over time revealed that it often exceeded the intensity of individual cofilin clusters measured on single filaments (Fig. 5D, Supp. Fig. 7). This observation indicates the presence of two colocalised cofilin clusters, one on each filament of the bundle.

In order to quantify the rate at which the second cofilin cluster is nucleated, we must take into account the other reactions taking place which together lead to the fragmentation of the 2-filament bundle. After a first cofilin cluster nucleation event on either filament of a 2-filament bundle, two different routes can lead to fragmentation. They are presented schematically in Fig. 5A. In the first route (‘route 1’), the first cofilin cluster severs the actin filament it is bound to, at a rate k_sev1_, before a cofilin cluster is nucleated on the other filament in the region facing the first cofilin cluster. Alternatively (‘route 2’), the second cofilin cluster is nucleated before the first cluster severs its actin filament. In route 2, the probability of nucleating a second cofilin cluster in the region where fascin has been excluded scales with the length of that region, which increases as the first cofilin cluster expands. The effective nucleation rate of the second cofilin cluster can thus be written as v_cof,1_ * t * k_nuc,2_, where v_cof,1_ is the growth velocity of the first cofilin cluster, k_nuc,2_ is the cofilin cluster nucleation rate per binding site in this fascin-free region for that cofilin concentration, and t is time. The competition between the two reactions, the severing of the first filament versus the nucleation of the second cluster, determines which route is followed by each 2-filament bundle.

In order to estimate k_nuc,2_ we sought to determine the relative importance of these two routes. One possibility is to identify severing events occurring on one of the two filaments within the bundle. The departure of one cofilin-saturated actin segment from the bundle, detectable as a drop in cofilin fluorescence is a good surrogate for the severing of a cofilin cluster (Supp. method text, Supp. Fig. 8).

We measured that ∼ 25% (± 8% std. dev., for a total of 217 bundles from 4 experiments) of the initial cofilin clusters fully severed and detached, leaving behind an unfragmented bundle with no detectable cofilin (Fig. 5 B,C). We interpreted these events as severing occurring before the nucleation of a second cofilin cluster, thus corresponding to route 1. At every moment, the chosen route depends on the rate of each reaction, and the rate of route 2 increases with time as the first cluster grows. Numerically fitting the cumulative time-distribution of events following route 1 (see Methods), allowed us to simultaneously determine both rates k_sev,1_ and k_nuc,2_ (Fig. 5C, Supp. Fig. 9). We obtained k_sev,1_ = 1.5 (± 1.2).10^-3^ s^-1^, which is comparable to the cofilin cluster severing rate measured on single filaments (rate k_sev,SF_ = 2.1 (± 0.3).10^-3^ s^-1^, Supp. Fig. 10). This indicates that the severing of cofilin clusters on 2-filament bundles is not significantly affected by fascin crosslinking. Remarkably, the nucleation rate of the second cofilin cluster, k_nuc,2_, is ∼ 8 times higher than the nucleation rate on single filaments (k_nuc,SF_), thus 48 times higher than the nucleation rate of the first cofilin clusters on 2-filament bundles (k_nuc,1_). This reveals the existence of a cofilin-driven ‘inter-filament cooperativity’, where the presence of a cofilin cluster on a filament strongly favors the nucleation of a cofilin cluster on the other filament of a 2-filament bundle. This inter-filament cooperativity probably arises from cofilin’s ability to locally twist actin filaments, which would be transmitted, at least partially, to the adjacent filament in the fascin-free uncrosslinked region. Such a mechanism would also relax the torsional stress induced by the cofilin cluster on the first filament, explaining why its severing rate is not enhanced as it would be if the filament was twist-constrained (Wioland *et al*, 2019b).

Following route 2, once there is a cofilin cluster on each filament of the bundle, each can sever at rate k_sev,2_ and the first severing event thus occurs with an apparent rate 2*k_sev,2_. This translates into a drop in cofilin fluorescence intensity to a plateau of similar amplitude as for individual cofilin clusters on single filaments (Fig. 5E). Fitting the cumulative time-distribution of these events yields k_sev,2_ ∼ k_sev,SF_ (Fig. 5F, see Methods). This indicates that the presence of two ‘side-by-side’ cofilin clusters on a bundle does not affect cofilin cluster severing rate. Ultimately, the severing of the remaining cofilin cluster occurs with a rate k_sev,final_ ∼ k_sev,SF_ (Fig. 5G), as expected for an individual cofilin cluster on a single filament.

Could the cofilin-driven inter-filament cooperativity we observed in 2-filament bundles be also at play in larger bundles? Although cofilin cluster nucleation events were rare on large bundles, we similarly could observe the nucleation of co-localized cofilin clusters in larger bundles, as revealed by cofilin fluorescence intensity (Supp. Fig. 11). On rare occasions, where large bundles fragmented completely (as quantified in Fig. 2E), we could observe multiple steps in the decrease of cofilin fluorescence intensity, indicating multiple cofilin cluster severing events. However, the complexity of large bundles, with a large number of events occuring in parallel, prevented us from conducting an analysis similar to the one we performed on 2-filament bundles. It thus seems that large bundles also exhibit inter-filament cooperativity for the nucleation of cofilin clusters.

### Cofilin twists actin filament bundles

In order to better understand the impact of cofilin on fascin-induced bundles, we used negative staining electron microscopy to observe their structure with a higher resolution. In the absence of cofilin, filaments in bundles are arranged in a parallel manner, as previously reported in vitro (Jansen *et al*, 2011). We were unable to observe 2-filament bundles in our samples, probably because freely diffusing filaments in solution tend to easily form large bundles in the presence of fascin. When exposed to cofilin, stretches of filaments of large bundles appeared braided, as if filaments were overtwisted and wrapped around each other (Fig. 6A).

From our kinetics analysis and electron microscopy observations, we propose a model that recapitulates cofilin binding and inter-filament cooperative cluster nucleation on fascin-induced 2-filament bundles (Fig. 6B). Initially, actin filaments in fascin-induced bundles are in conformations that are less favorable for cofilin binding than isolated actin filaments. Once a cofilin cluster has nucleated, its expansion locally triggers fascin unbinding and prevents fascin from rebinding. Cofilin-induced increase of filament helicity causes a local twisting of the entire bundle, thereby changing the helicity of the fascin-free filament region, facing the cofilin cluster. In this region, the increase in helicity enhances cofilin affinity, and thus locally favors the nucleation of a cofilin cluster.

### Nucleation of the first cofilin cluster is the limiting factor in the fragmentation of bundles

To recapitulate the impact of fascin-induced bundling on filament disassembly by cofilin, we performed Gillespie simulations (Fig. 7A), integrating all the individual reaction rates that we measured experimentally in microfluidics assays (Figs. 3-5). First, we compared the fraction of unsevered 2-filament bundles and unsevered single filaments, as a function of time (n = 200 simulated 5-µm long segments for each population). Without any free parameter, the simulated curves are in good agreement with the experimental data (Fig. 7B). In the simulations, all 2-filament bundle fragmentation events that occurred before 1500 seconds were caused by cooperatively-nucleated overlapping cofilin clusters (route 2 in Fig. 5), as experimentally observed. This indicates that, over these time scales, inter-filament cooperativity is the dominant pathway leading to bundle fragmentation, and that the nucleation of a new cluster where one filament has already severed (route 1 in Fig. 5) is extremely rare, explaining why we never observed these final steps in our experiments.

In order to assess the impact of the different steps leading to bundle fragmentation, we modified the reaction rates in our numerical model. We focused on the two key effects of fascin-induced bundling: the hindered nucleation of the first cofilin cluster (low k_nuc,1_), which delays the fragmentation of the bundle, and inter-filament cooperativity for the nucleation of the second cofilin cluster (high k_nuc,2_), which favors the fragmentation of the bundle. We first simulated the situation where cofilin cluster nucleation is unaffected by fascin crosslinking (i.e. k_nuc,1_ = k_nuc,2_ = k_nuc,SF_), keeping all the other rates unchanged. In this situation, there is no hindrance of the first cluster nucleation and no inter-filament cooperativity. This resulted in a ∼2-fold faster bundle fragmentation, compared to what we observed experimentally (Supp. Fig. 12). We then simulated the situation where the first cluster nucleation is hindered, but where there is no inter-filament cooperativity (i.e. k_nuc,1_ = k_nuc,2_ = 1/6 k_nuc,SF_). In this situation, bundle fragmentation is ∼2-fold slower than in our experiments (Supp. Fig. 12). These two alternative simulated scenarios indicate that both the delayed nucleation of the initial cofilin clusters and the inter-filament cooperativity of the following cofilin clusters strongly impact the rate at which 2-filament bundles are fragmented by cofilin.

### Twist-constrained bundle fragmentation

When single actin filaments are constrained in twist, by being attached to fixed anchors at each end, cofilin severing is dramatically increased due to the inability of filaments to relax cofilin-induced torsional stress (Wioland *et al*, 2019b). In cells, filopodia making contacts with the extracellular matrix or other cells could constrain the twist of fascin-induced actin filament bundles (Jacquemet *et al*, 2015). We thus sought to quantify the cofilin-induced fragmentation of twist-constrained fascin-induced 2-filament bundles. To do so, the downstream part of filament bundles were anchored to the glass surface, using biotin-streptavidin linkages (Fig. 7C). We observed that twist-constrained bundles fragmented significantly faster than unanchored ones (Supp. Fig. 13). We observed that the nucleation rate of cofilin clusters was similar for both twist-constrained and twist-unconstrained fascin bundles (Supp. Fig. 14), in agreement with observations on single actin filaments (Wioland *et al*, 2019b).

The rapid fragmentation of twist-constrained 2-filament bundles prevented us from directly quantifying the nucleation rate of the subsequent cofilin clusters that overlapped with the initial ones. In order to assess inter-filament cooperativity for twist-constrained bundles, we performed numerical simulations where the growth of cofilin clusters creates a torque and increases the severing rates of cofilin clusters. As above, numerical simulations were performed using parameters determined experimentally (Fig. 2-5), and with an exponentially increasing cofilin-induced severing rate as torque accumulates, as previously characterized for single twist-constrained filaments (Wioland *et al*, 2019b) (Fig. 7C). In simulations with an inter-filament cooperative cofilin nucleation rate of the same amplitude as the one determined for unconstrained bundles (k_nuc,2_ = 8 k_nuc,SF_), simulated twist-constrained bundles fragmented appreciably faster than experimentally observed (Supp. Fig 15). Simulations performed without inter-filament cooperative nucleation (k_nuc,2_ = k_nuc,SF_) appear in better agreement with our experimental observations (Fig. 7C). In this latter case, 75 % of first cofilin clusters severed before a second overlapping cofilin cluster could be nucleated.

Overall, these results can be consistently interpreted in the frame of our model (Figs 5A and 6B). They indicate that constraining the twist of fascin-induced filament bundles changes mainly two of the steps that lead to their fragmentation by cofilin. First, constraining the twist prevents the supercoiling of the bundle, which requires its ability to rotate freely (Fig 6B) and thus abolishes the inter-filament cooperative nucleation of the second cofilin clusters. Second, constraining the twist prevents the relaxation of the mechanical torque induced by the first cofilin cluster and accelerates its severing.

## Discussion

In this study, we reveal that actin filaments are severed more slowly by cofilin when they are bundled by fascin, and we decipher the molecular details underlying this phenomenon. In particular, we show that bundle fragmentation is slower than what would occur if each filament in the bundle behaved as an individual filament and fragmentation of the bundle would occur upon the colocalization of independent severing events on each filament. Strikingly, this is primarily due to the slower nucleation of cofilin clusters on fascin-induced bundles than on single filaments. In addition, one could have expected fascin to constrain filament twist, leading to faster severing of filaments by cofilin (Wioland *et al*, 2019b; Breitsprecher *et al*, 2011). We observed that this is not the case for 2-filament bundles, and appears unlikely for larger bundles as well. On the contrary, our observations indicate that a local change of filament helicity is induced by cofilin binding and that it is transmitted to the adjacent filaments in the bundle. To our knowledge, this is the first time such an inter-filament cooperativity between actin filaments is ever reported. Together with the exclusion of fascin from cofilin-decorated regions, this transmission of the local change of filament helicity strongly favors the nucleation of cofilin clusters on adjacent filaments. This inter-filament cooperativity mechanism leads to the co-localization of cofilin clusters, and permits bundle fragmentation faster than what would have been observed if the nucleation of cofilin clusters on adjacent filaments were purely random. Overall, these observations show that fascin crosslinking strongly impacts cofilin activity, with two opposing effects: First, it hinders the nucleation of the first cofilin clusters, on cofilin-free regions of the bundles, and, second, it greatly accelerates the nucleation of additional cofilin clusters on bare filaments, adjacent to an already existing cluster. Globally, the first effect dominates, the second one does not compensate, and fascin-induced bundles are protected from severing. This is especially the case for large bundles.

Our study provides new molecular insights into the initial binding of cofilin on intact fascin-induced bundles and on the importance of bundle size. Using pyrene-actin bulk experiments, Breitsprecher and colleagues previously observed a weaker cofilin binding to fascin-induced filament bundles (Breitsprecher *et al*, 2011). However, based on other observations, using various experimental approaches, they proposed that fascin acted as anchors along filaments and prevented cofilin from changing filament helicity, and thus accelerated cofilin severing of filaments in the bundles. This proposed mechanism is somewhat similar to what we reported for artificially twist-constrained single actin filaments (Wioland *et al*, 2019b). Here, we reveal that fascin is locally excluded from cofilin-decorated segments. This exclusion of fascin has important consequences for the disassembly of the bundles as it allows cofilin to locally change the helicity of filaments, thereby preventing it from generating a torque that would accelerate severing. In the case where filament bundles are anchored to the surface of the chamber, twist is constrained, cofilin-induced inter-filament cooperativity is suppressed and bundles fragment rapidly due to accelerated cofilin severing (Wioland *et al*, 2019b).

Why is cofilin binding hindered on bundles? Fascin binds in between two bundled filaments every 36 nm (Aramaki *et al*, 2016; Jansen *et al*, 2011). Many actin subunits are therefore available for cofilin to bind, between two consecutive bound fascins. Consequently, direct steric competition between fascin and cofilin cannot fully account for the reduction in cofilin binding on the filaments of a bundle. However, fascin bundling moderately decreases the helicity of filaments, by ∼1° (Shin *et al*, 2009). One hypothesis explaining the slower cofilin binding could therefore be that, due to fascin bundling, filaments are ‘trapped’ in conformations that are less favorable for cofilin binding. This interpretation complies with the notion of ‘filament breathing’, where thermal fluctuations of the filament conformation (subunit conformations, inter-subunit interfaces, local filament curvature and twist, etc.) strongly modulate the binding of regulatory proteins (Galkin *et al*, 2010; Schramm *et al*, 2019; Reynolds *et al*, 2022). Here, fascin would decrease the relative time spent by actin filaments in conformations that are preferential for cofilin to bind. We have previously shown that cofilin binding to twist-constrained actin filaments is unaffected (Wioland *et al*, 2019b). We therefore argue that reducing the fluctuations of actin filaments has probably a stronger impact on cofilin binding than shifting the average helicity imposed by ∼1° (Jégou & Romet-Lemonne, 2020).

Another important aspect of fascin-induced bundling is filament packing. As opposed to larger bundlers (e.g. alpha-actinin), fascin tightly packs filaments, with an inter-filament distance of only ∼ 6 nm (Jansen *et al*, 2011). The accessibility of filaments at the core of bundles for cofilin that are diffusing in solution could potentially be partially hindered. Actually, to reach the core, cofilin has to diffuse through a medium composed of absorbing obstacles (i.e. the actin filaments of the outer shell of bundles)(Saxton, 1994), so cofilin could be locally depleted (Manhart *et al*, 2019), and filaments at the core of bundles be ‘protected’ from cofilin. We indeed show that cofilin binds less efficiently to larger bundles (Fig. 3). For larger bundles, one would have to wait for filaments in the periphery to first be targeted by cofilin to make filaments of the inner core accessible to cofilin. In conclusion, fascin-induced bundling could thus hinder cofilin binding through various mechanisms, and their relative contributions are difficult to disentangle. In cells, fascin has been identified as the main crosslinker of actin filaments in filopodia (Vignjevic *et al*, 2006; Aramaki *et al*, 2016). Our results indicate that fascin-induced bundles in growing filopodia are potentially protected from cofilin in the cytoplasm, and fascin gets excluded locally upon cofilin binding. Indeed, such a phenotype has been recently reported in filopodia of neuronal growth cones, thanks to EM tomography. In one study, some filaments within filopodia displayed segments with a shorter pitch, and appeared disconnected from the hexagonally packed filament bundles (Atherton *et al*, 2022). Another study further revealed regions of filament bundles decorated by cofilin within filopodia using live fluorescence microscopy, in addition to EM observations (Hylton *et al*, 2022). These regions coincided with filopodia kinks or wavy shapes. These deformation of filopodia are also reminiscent of the report of filopodia rotating in a clockwise fashion (Tamada, 2019; Leijnse *et al*, 2022), which could thus be attributable to cofilin inter-filament cooperativity inducing actin filament bundle rotation. Rotation of filopodia in cells has been proposed to be an essential aspect of the emergence of the left-right asymmetry during brain development (Tamada, 2019) and could thus rely on cofilin. Lastly, besides filopodia, fascin was also implicated in the maturation of stress fibers, locally protecting filaments connected to focal adhesions from cofilin disassembly (Elkhatib *et al*, 2014). This illustrates further how fascin influences the stability of various actin structures in cells.

Is bundle complete fragmentation happening in cells? While cofilin probably does sever filaments constituting bundles, the high concentration of profilin-actin in cells (Funk *et al*, 2019) may allow for the regrowth of severed filament barbed ends along bundles, at least in cases where filament barbed ends are not saturated by cofilin (Wioland *et al*, 2017). Newly growing barbed ends may then be crosslinked ‘back’ into the bundle, thanks to the free fascin available in the cytosol (Vignjevic *et al*, 2006), and this would be a possible self-repairing mechanism. However, these newly generated barbed ends could just as easily be rapidly capped by capping proteins CP which is also present in filopodia (Sinnar *et al*, 2014; Edwards *et al*, 2014). The final outcome would thus be balanced by the relative abundance of these regulatory proteins and is hard to predict.

Apart from filament disassembly, we envision that cofilin could potentially play a so far unanticipated role, by inducing the rotation of parts of actin networks. Here, we reveal that geometrical organization is a factor that modulates cofilin activity, but also that cofilin can change the shape of actin networks. This may also apply to networks made by other crosslinking proteins. For example, alpha-actinin (Christensen *et al*, 2017), which is larger and more flexible than fascin, may differentially impact cofilin ability to reorganize actin bundles. Interestingly, in fibroblasts, ADF/cofilin has been recently identified to drive the establishment of chiral radial stress fibers (Tee *et al*, 2023), for which alpha-actinin is thought to be the main crosslinker. Our results thus open new perspectives on how filament crosslinking integrates in a global scheme that tune actin network turnover and mechanical properties.

## METHODS

### Biochemistry

#### Protein purification

α-skeletal muscle actin was purified from rabbit muscle acetone powder following the protocol described in (Wioland et al, 2017), based on the original protocol from (Spudich and Watt, 1971).

Spectrin-actin seeds were purified from human erythrocytes as described in (Wioland et al, 2017), based on the original protocol by (Casella et al, 1986).

Human fascin-1 (Uniprot: Q16658): 6xHis-fascin1 was purified as described previously in (Suzuki et al, 2020).

Mouse cofilin-1 (Uniprot: P18760): fluorescent fusion protein 6xHis-eGFP-cofilin-1 and 6xHis-mCherry-cofilin-1 were purified as described previously in (Kremneva et al, 2014).

#### Protein labelling

Actin was fluorescently labeled on accessible surface lysines of F-actin, using Alexa-488 succinimidyl ester (Life Technologies). To minimize effects from the fluorophore we used a labeling fraction of 10 % for both microfluidics and open chamber assays.

Actin was similarly labeled with biotin, using biotin succinimidyl ester (Life Technologies).

Fascin was labeled on surface cysteines using Alexa-568 maleimide (Life Technologies), leading to a labeling fraction of ∼ 180 %.

Cofilin-1 was fused with mCherry or eGFP at their N-terminus (Kremneva et al, 2014). We systematically used 100 % labeled cofilin-1, for which activity has been verified previously (Wioland et al. 2017).

#### Fluorescence Microscopy

For both ‘open chambers’ and microfluidics assays, we took care to minimize the impact of light exposure that impacts cofilin binding to actin filaments.

We observed day to day variations in the activity of cofilin. To minimize this limitation, we took advantage of the microfluidics experiments: we exposed single filaments and filament bundles that are side-by-side at the surface of the chamber to the same cofilin conditions, in order to derive fold-change between cofilin activities on single filaments and bundles.

#### Buffers

All experiments were performed in F-buffer: 5 mM Tris HCl pH 7.4, 50 mM KCl, 1 mM MgCl_2_, 0.2 mM EGTA, 0.2 mM ATP, 10 mM DTT and 1 mM DABCO. The concentrations of DTT and DABCO were chosen to limit light-induced artifacts. Buffers were supplemented with 0.2% methylcellulose (4000 cP at 2%, Sigma) for open chamber assays to keep filaments in the vicinity of the glass bottom and image them using TIRF microscopy (with a laser penetration depth ∼ 80 nm).

#### Cofilin binding assay in ‘open chambers’

Experiments were conducted in chambers made by assembling two 22×40 mm #1.5 coverslips, spaced by a melted parafilm. The surface was incubated with 2 pM spectrin-actin seeds for 5 minutes, then passivated with bovine serum albumin (BSA, 50 mg/mL, for 5 minutes). Actin filaments were elongated from surface-anchored spectrin-actin seeds, using 10% labeled Alexa488-actin in F-buffer, supplemented with 0.2% methylcellulose (4000 cP at 2%, Sigma). They are then aged to become fully ADP-actin filaments by exposing them to 0.15 µM Alexa488-actin for 15 minutes, in the presence or absence of 200 nM fascin. Filaments are then exposed to 80 nM mCherry-cofilin1 and 200 nM fascin. Note that the cofilin concentration used in open chambers is substantially lower than in microfluidics experiments, due to the propensity of methylcellulose to increase protein concentration close to the glass bottom.

#### Bundles and single filaments in microfluidics

Microfluidics experiments were done with Poly-Dimethyl-Siloxane (PDMS, Sylgard) chambers based on the original protocol from (Jégou *et al*, 2011), and described in detail in (Wioland *et al*, 2022). Briefly, glass coverslips previously were cleaned in sequential ultrasound baths of 2% Hellmanex, 2M KOH, pure water and ethanol, each for 30 minutes and extensive rinsing in pure water between each step. A PDMS cross-shaped chamber with 3 inlets (inner main channel dimensions of 20 μm in height, 800 μm in width and ∼1 cm in length) was mounted onto a cleaned glass coverslip, both previously plasma-activated for 30 seconds to allow them to bind tightly to each other. The microfluidics chamber was connected to solution reservoirs by blue PEEK tubings. Pressure in the reservoirs was controlled and flow rates monitored using microfluidic devices MFCS-EZ and Flow Units (from Fluigent). The temperature was controlled and set to 25°C (using an objective-collar heater from Oko-lab).

We used spectrin-actin seeds to anchor filaments by their pointed-end to the microfluidics coverslip surface: the surface was incubated with 2 pM spectrin-actin seeds for 5 minutes, then passivated with bovine serum albumin (BSA, 50 mg/mL, for 5 minutes). Filaments were then grown using a solution of 1 μM 10% Alexa-488-G-actin in F-buffer. Filaments are finally aged for at least 15 minutes with G-actin at critical concentration (0.15 μM, 10% Alexa-488 fluorescently labeled), to ensure that the actin is > 99.9% in ADP-state, in the presence or absence of 200 nM fascin. We verified that fascin bundling did not significantly slow down Pi-release during the aging process (Supp. Fig. 2).

#### Twist-constrained filaments and bundles

To constrain the twist of single filaments and bundles (Fig. 7), filaments were first grown from surface-anchored spectrin-actin seeds in a microfluidics chamber previously passivated by 50:1 BSA:biotin-BSA (0.5 mg/mL in F-buffer, for 5 minutes). Filaments were sequentially elongated, first using 1 μM Alexa-488-G-actin to grow ∼ 10 µm-long segments, then using 0.5 µM 1 % biotin-labeled actin for 1 minute to generate a ∼ 2 µm-long biotinylated segment at their barbed end. Filaments were then aged for 15 minutes, with Alexa-488-G-actin at critical concentration (0.15 μM), to ensure that the actin is in > 99.9% ADP-state, in the presence of 200 nM fascin. Bundles were subsequently exposed to neutravidin (3 µg/mL) in the presence of 200 nM fascin and 0.15 µM actin, for 5 minutes, to anchor their distal end to the biotinylated surface. Finally, bundles were exposed to 200 nM mCherry-cofilin-1, 200 nM fascin, and 0.15 µM Alexa488(10%)-G-actin, to quantify the nucleation rate of cofilin clusters and the fragmentation of twist-constrained bundles and single filaments.

#### Formation of larger bundles

To form fascin-induced bundles composed of more than 2 filaments, filaments were grown from non-specifically adsorbed 1 µm in diameter glass beads. Beads were previously functionalized with spectrin-actin seeds (100 pM in F-buffer for 5 minutes, rinsed twice in F-buffer). The coverslip surface was then passivated with BSA (50 mg/mL) for at least 5 minutes. The number of filaments per bundle was quantified using actin fluorescence prior to cofilin exposure (Supp. Fig 4A), using the fluorescence of single filaments that grew from the surface as a reference.

#### Image acquisition

Experimental chambers were positioned on a Nikon TiE inverted microscope, equipped with a 60x oil-immersion objective. In microfluidics, we systematically used the epifluorescence illumination mode to avoid fluorescence variation due to filament or bundle height in the evanescent/TIRF illumination mode. For open chamber assays, methylcellulose strongly constrains filament and bundle height to 50 nm above the glass surface, and TIRF illumination was used. 100 mW tunable lasers (iLAS2, Gataca Systems) were used for TIRF illumination, or a 120W Xcite Exacte lamp (Lumen Dynamics) for epifluorescence imaging. TiE microscopes were controlled either by Metamorph or ImageJ/Micromanager (Edelstein et al, 2014) softwares. Images were acquired by an Evolve EMCCD (Photometrics) or an sCMOS Orca-Flash4.0 V3.0 (Hamamatsu) camera.

#### Negative Staining Electron Microscopy

Three microlitres of sample were deposited on a 400 mesh copper grid with a conventional glow discharge activated carbon film. After 1 minute, the sample was briefly drained by Whatman paper filter and the grid was covered with a drop of 1% (w/v) aqueous uranyl acetate for 30 seconds and then dried. The grids were examined at 120kV with TEM (Tecnai 12, Thermo Fischer Scientific) equipped with a 4K CDD camera (Oneview, Gatan).

#### Data analysis

For both single filaments and bundles, measurements were performed on 5 µm-long segments. For single filaments, we excluded the region that extends 2 pixels away from the seed position due to the sharp bending of the filament. For bundles, as they are formed from filaments elongated from surface-anchored seeds which might be microns away from each other, only the central regions for which the actin fluorescence intensity clearly indicates that 2 filaments are bundled together were analyzed.

During the course of an experiment, severing events often occur at the seed location due to the sharp bent of the filament (Wioland *et al*, 2019b). All these events do not correspond to regular severing events and were taken into account as ‘censoring’ events. We use the Kaplan-Meier method to estimate survival fractions and confidence intervals (Kaplan, 1958), which are implemented in the ‘lifelines’ package in python.

Cofilin cluster nucleation on ADP-actin filaments or bundles were quantified by manually detecting the appearance of cofilin clusters giving rise to detectable growing clusters. The time-dependent cumulative distribution of the fraction of actin segments with at least one nucleated cofilin cluster is fitted by a single exponential to obtain the cofilin cluster nucleation rate per actin binding site per second.

Cofilin cluster growth rates were obtained using kymographs, and by manually measuring the width of cofilin clusters for at least 3 different time points.

Cofilin cluster severing events and filament/bundle fragmentations were manually tracked and listed as time points to build ‘survival’ curves and fitted as follows:

- The population of isolated cofilin clusters on 2-filament bundles that severed before a second cofilin cluster could be detected (Fig. 5) was fitted using a chi-square minimization procedure (‘curve_fit’ from the scipy package in python), by simulating, using the Euler method, the outcome of a population of 200 cofilin clusters that grow at a rate v_growth_ and can either sever with a rate k_sev,1_ or cooperatively nucleate a new cofilin cluster on the adjacent filament in the region that overlaps with the initial cofilin cluster, at a rate k_nuc,2_ * t * v_growth_, where k_nuc,2_ is the rate of nucleation of the second cofilin cluster. The least-square fitting procedure gives the most probable values for the two free parameters, k_sev,1_ and k_nuc,2_.
- The survival fraction of two-overlapping cofilin clusters on 2-filament bundles (Fig. 5) was fitted using a chi-square minimization procedure (‘curve_fit’ from the scipy package in python), by simulating, using the Euler method, the outcome of a population of 200 cofilin clusters that grow at a rate v_growth_, allow the nucleation, at a rate k_nuc,2_, of a second cofilin cluster that overlaps with the first one and grows at the similar rate v_growth_, and the two cofilin clusters sever at an effective rate 2 x k_sev,2_. The least-square fitting procedure gives the most probable value for the only free parameter, k_sev,2_.
- The survival fraction of the isolated cofilin clusters that arise from an initial population of two overlapping cofilin clusters on 2-filament bundles, and where one of them has already severed (Fig. 5), is fitted by a single exponential decay function with a decay rate k_sev,final_.
- Single filament and bundle fragmentation were fitted by a single exponential decay function in order to derive a fold difference in the rate at which fragmentation occurs for both populations (Fig. 2).

### Numerical Simulations

We performed Gillespie numerical simulations to simulate cofilin activity on single filaments (1 segment) or 2-filament bundles (2 segments), each segment being 5 µm long (1818 actin subunits in length). Absolute parameters used in the simulations are reported in supplementary table 1. We applied the following rules:

- a cofilin cluster nucleates at a rate:

_-_ on bare actin subunits of filaments of 2-filaments bundles: k_nuc,bundle_
_-_ on bare actin subunits of single filaments: k_nuc,SF_ = 6 x k_nuc,bundle_
- on actin subunits that face a cofilin-saturated actin region on the adjacent filament of 2-filament bundles: k_nuc,bundleEnhanced_ = 48 x k_nuc,bundle_
_-_ on actin subunits that face a cofilin-occupied actin region on the adjacent filament of 2-filament bundles, and which at least one of the boundaries of this latter cluster has severed: k_nuc,SF_
- a cofilin cluster grows symmetrically towards both the pointed and barbed ends, at a rate v_growth_, in all cases.
- a cofilin cluster severs at a rate k_sev_, in all cases, 80% (resp. 20%) of the time towards the pointed end (resp. barbed end), as previously reported in (Wioland *et al*, 2017).
- for twist-constrained single filaments or bundles, a cofilin cluster severs at a rate that is exponentially increased as a function of the cofilin-induced torque, using k_sev,torque_ = k_sev_ x exp(α.Γ / k_B_.T), where α is a constant whose value was set to 5, to match the previously best reported value in (Wioland *et al*, 2019b), Γ is the applied torque (in pN.nm) and k_B_.T is the product of the Boltzmann constant and the temperature (4.1 pN.nm, at 25°C). The torque was computed as in (Wioland *et al*, 2019b), using Γ = ν . dθ / dL . C_a_ . C_c_ / (ν.C_a_ + (1-ν). C_c_), with ν the cofilin density along the segment, dθ = 4.7° the rotation along the long filament axis induce by the binding of an additional cofilin molecule, dL = 2.7 nm the length of an actin subunits in the filament, and C_a_ (resp. C_c_) the torsional rigidity of a bare (resp. cofilin-decorated) single actin filament. We used previously reported values from (Prochniewicz *et al*, 2005) C_a_ = 2.3 . 10^3^ pN.nm^2^/rad, and C_c_ = 0.13 . 10^3^ pN.nm^2^/rad.
- The simulation is halted and the time of the event is recorded for analysis, if:

- for both single filaments and 2-filament bundles: they are fully decorated by cofilin and no fragmentation has occurred, or the simulation time has reached t_max_ = 3000 seconds (8000 s, for Supp. Fig. 12).
- for single filaments: the filament is fragmented, i.e. upon the first cofilin cluster severing event.
- for 2-filament bundles: when two (partially) severed cofilin-decorated regions, one on each filament, are co-localized. In this case, fascin cannot crosslink all the actin segments together.

### Softwares

All measurements (length, distance, fluorescence intensity, severing time,…) were performed manually on Fiji/ImageJ. Data analysis and statistical significance tests were done using python (with numpy, scipy, panda and lifelines packages). Gillespie numerical simulations were performed using python.

### Statistical significance

Comparison of the survival distributions of two samples were done using the p-value from the log-rank test, using the lifelines package in python.

The sample data means were compared using the Welch’s paired two-samples t-test in order to derive a p-value, using the ‘ttest_rel’ function from the scipy package in python. Superplots were generated using SuperPlotsOfData – a web app for the transparent display and quantitative comparison of continuous data from different conditions (Goedhart, 2021), available online at https://huygens.science.uva.nl/SuperPlotsOfData/.

## Supporting information

Supplementary Informations

## ACKNOWLEDGEMENTS

JC, LL and AJ were supported by the European Research Council (ERC) under the European Union’s Horizon 2020 research and innovation program (grant agreement StG-679116 to AJ). JC was also supported by the Labex WhoAmI? of Université Paris Cité. GRL and HW were supported by Agence Nationale de la Recherche (grant RedoxActin to GRL). We acknowledge the ImagoSeine core facility of the Institut Jacques Monod, member of the France BioImaging infrastructure (ANR-10-INBS-04) and GIS-IBiSA.

For the purpose of Open Access, the authors have applied a CC BY public copyright license to any Author Accepted Manuscript version arising from this submission.

## REFERENCES

Adams JC & Schwartz MA (2000) Stimulation of fascin spikes by thrombospondin-1 is mediated by the GTPases Rac and Cdc42. J Cell Biol 150: 807–822

Aramaki S, Mayanagi K, Jin M, Aoyama K & Yasunaga T (2016) Filopodia Formation by Cross-linking of F-actin with Fascin in Two Different Binding Manners. Cytoskeleton

Aratyn YS, Schaus TE, Taylor EW & Borisy GG (2007) Intrinsic dynamic behavior of fascin in filopodia. Mol Biol Cell 18: 3928–3940

Atherton J, Stouffer M, Francis F & Moores CA (2022) Visualising the cytoskeletal machinery in neuronal growth cones using cryo-electron tomography. J Cell Sci 135

Banerjee S, Gardel ML & Schwarz US (2020) The Actin Cytoskeleton as an Active Adaptive Material. Annu Rev Condens Matter Phys

Blanchoin L, Boujemaa-Paterski R, Sykes C & Plastino J (2014) Actin Dynamics, Architecture, and Mechanics in Cell Motility. Physiol Rev 94: 235–263

Blanchoin L & Pollard TD (1999) Mechanism of interaction of Acanthamoeba actophorin (ADF/Cofilin) with actin filaments. J Biol Chem 274: 15538–15546

Breitsprecher D, Koestler SA, Chizhov I, Nemethova M, Mueller J, Goode BL, Small JV, Rottner K & Faix J (2011) Cofilin cooperates with fascin to disassemble filopodial actin filaments. J Cell Sci 124: 3305–3318

Campbell JJ & Knight MM (2007) An improved confocal FRAP technique for the measurement of long-term actin dynamics in individual stress fibers. Microsc Res Tech 70: 1034–1040

Carlier MF, Laurent V, Santolini J, Melki R, Didry D, Xia GX, Hong Y, Chua NH & Pantaloni D (1997) Actin depolymerizing factor (ADF/cofilin) enhances the rate of filament turnover: implication in actin-based motility. J Cell Biol 136: 1307–1322

Chalut KJ & Paluch EK (2016) The Actin Cortex: A Bridge between Cell Shape and Function. Dev Cell 38: 571–573

Christensen JR, Hocky GM, Homa KE, Morganthaler AN, Hitchcock-DeGregori SE, Voth GA & Kovar DR (2017) Competition between Tropomyosin, Fimbrin, and ADF/Cofilin drives their sorting to distinct actin filament networks. Elife 6

Claessens MMAE, Bathe M, Frey E & Bausch AR (2006) Actin-binding proteins sensitively mediate F-actin bundle stiffness. Nat Mater 5: 748–753

Claessens MMAE, Semmrich C, Ramos L & Bausch AR (2008) Helical twist controls the thickness of F-actin bundles. Proceedings of the National Academy of Sciences 105: 8819–8822

Damiano-Guercio J, Kurzawa L, Mueller J, Dimchev G, Schaks M, Nemethova M, Pokrant T, Brühmann S, Linkner J, Blanchoin L, et al (2020) Loss of Ena/VASP interferes with lamellipodium architecture, motility and integrin-dependent adhesion. Elife 9

De La Cruz EM (2005) Cofilin binding to muscle and non-muscle actin filaments: isoform-dependent cooperative interactions. J Mol Biol 346: 557–564

Edwards M, Zwolak A, Schafer DA, Sept D, Dominguez R & Cooper JA (2014) Capping protein regulators fine-tune actin assembly dynamics. Nat Rev Mol Cell Biol: 1–13

Edwards RA, Herrera-Sosa H, Otto J & Bryan J (1995) Cloning and expression of a murine fascin homolog from mouse brain. J Biol Chem 270: 10764–10770

Elkhatib N, Neu MB, Zensen C, Schmoller KM, Louvard D, Bausch AR, Betz T & Vignjevic DM (2014) Fascin plays a role in stress fiber organization and focal adhesion disassembly. Curr Biol 24: 1492–1499

Faix J, Breitsprecher D, Stradal TEB & Rottner K (2009) Filopodia: Complex models for simple rods. Int J Biochem Cell Biol 41: 1656–1664

Freedman SL, Suarez C, Winkelman JD, Kovar DR, Voth GA, Dinner AR & Hocky GM (2019) Mechanical and kinetic factors drive sorting of F-actin cross-linkers on bundles. Proc Natl Acad Sci U S A 116: 16192–16197

Fritzsche M, Lewalle A, Duke T, Kruse K & Charras G (2013) Analysis of turnover dynamics of the submembranous actin cortex. Mol Biol Cell 24: 757–767

Funk J, Merino F, Venkova L, Heydenreich L, Kierfeld J, Vargas P, Raunser S, Piel M & Bieling P (2019) Profilin and formin constitute a pacemaker system for robust actin filament growth. Elife 8: :e50963

Galkin VE, Orlova A, Lukoyanova N, Wriggers W & Egelman EH (2001) Actin Depolymerizing Factor Stabilizes an Existing State of F-Actin and Can Change the Tilt of F-Actin Subunits. Journal of Cell Biology 153: 75–86 doi:10.1083/jcb.153.1.75 [PREPRINT]

Galkin VE, Orlova A, Schröder GF & Egelman EH (2010) Structural polymorphism in F-actin. Nat Struct Mol Biol 17: 1318–1323

Gallop JL (2019) Filopodia and their links with membrane traffic and cell adhesion. Semin Cell Dev Biol

Goedhart J (2021) SuperPlotsOfData-a web app for the transparent display and quantitative comparison of continuous data from different conditions. Mol Biol Cell 32: 470–474

Gressin L, Guillotin A, Guérin C, Blanchoin L & Michelot A (2015) Architecture dependence of actin filament network disassembly. Curr Biol 25: 1437–1447

Hotulainen P, Paunola E, Vartiainen MK & Lappalainen P (2005) Actin-depolymerizing factor and cofilin-1 play overlapping roles in promoting rapid F-actin depolymerization in mammalian nonmuscle cells. Mol Biol Cell 16: 649–664

Huehn AR, Bibeau JP, Schramm AC, Cao W, De La Cruz EM & Sindelar CV (2020) Structures of cofilin-induced structural changes reveal local and asymmetric perturbations of actin filaments. Proc Natl Acad Sci U S A

Hylton RK, Heebner JE, Grillo MA & Swulius MT (2022) Cofilactin filaments regulate filopodial structure and dynamics in neuronal growth cones. Nat Commun 13: 2439

Ishikawa R, Sakamoto T, Ando T, Higashi-Fujime S & Kohama K (2003) Polarized actin bundles formed by human fascin-1: their sliding and disassembly on myosin II and myosin V in vitro. J Neurochem 87: 676–685

Jacquemet G, Hamidi H & Ivaska J (2015) Filopodia in cell adhesion, 3D migration and cancer cell invasion. Curr Opin Cell Biol 36: 23–31

Jacquemet G, Stubb A, Saup R, Miihkinen M, Kremneva E, Hamidi H & Ivaska J (2019) Filopodome Mapping Identifies p130Cas as a Mechanosensitive Regulator of Filopodia Stability. Curr Biol 29: 202–216.e7

Jansen S, Collins A, Yang C, Rebowski G, Svitkina T & Dominguez R (2011) Mechanism of actin filament bundling by fascin. J Biol Chem 286: 30087–30096

Jégou A, Niedermayer T, Orbán J, Didry D, Lipowsky R, Carlier M-F & Romet-Lemonne G (2011) Individual actin filaments in a microfluidic flow reveal the mechanism of ATP hydrolysis and give insight into the properties of profilin. PLoS Biol 9: e1001161

Jégou A & Romet-Lemonne G (2020) Mechanically tuning actin filaments to modulate the action of actin-binding proteins. Curr Opin Cell Biol 68: 72–80

Lai FPL, Szczodrak M, Block J, Faix J, Breitsprecher D, Mannherz HG, Stradal TEB, Dunn GA, Small JV & Rottner K (2008) Arp2/3 complex interactions and actin network turnover in lamellipodia. EMBO J 27: 982–992

Lappalainen P, Kotila T, Jégou A & Romet-Lemonne G (2022) Biochemical and mechanical regulation of actin dynamics. Nat Rev Mol Cell Biol

Leijnse N, Barooji YF, Arastoo MR, Sønder SL, Verhagen B, Wullkopf L, Erler JT, Semsey S, Nylandsted J, Oddershede LB, et al (2022) Filopodia rotate and coil by actively generating twist in their actin shaft. Nat Commun 13: 1636

Lieleg O, Claessens MMA & Bausch AR (2010) Structure and dynamics of cross-linked actin networks. Soft Matter 6: 218–225

Maciver SK, Zot HG & Pollard TD (1991) Characterization of actin filament severing by actophorin from Acanthamoeba castellanii. J Cell Biol 115: 1611–1620

Manhart A, Icheva TA, Guerin C, Klar T, Boujemaa-Paterski R, Thery M, Blanchoin L & Mogilner A (2019) Quantitative regulation of the dynamic steady state of actin networks. Elife 8

Ma R & Berro J (2018) Structural organization and energy storage in crosslinked actin assemblies. PLoS Comput Biol 14: e1006150

McCullough BR, Blanchoin L, Martiel J-L & De la Cruz EM (2008) Cofilin increases the bending flexibility of actin filaments: implications for severing and cell mechanics. J Mol Biol 381: 550–558

McGough A, Pope B, Chiu W & Weeds A (1997) Cofilin changes the twist of F-actin: implications for actin filament dynamics and cellular function. J Cell Biol 138: 771–781

Prochniewicz E, Janson N, Thomas DD & De la Cruz EM (2005) Cofilin increases the torsional flexibility and dynamics of actin filaments. J Mol Biol 353: 990–1000

Rajan S, Kudryashov DS & Reisler E (2023) Actin Bundles Dynamics and Architecture. Biomolecules 13: 450

Reynolds MJ, Hachicho C, Carl AG, Gong R & Alushin GM (2022) Bending forces and nucleotide state jointly regulate F-actin structure. Nature: 1–7

Saito T, Matsunaga D & Deguchi S (2022) Long-term molecular turnover of actin stress fibers revealed by advection-reaction analysis in fluorescence recovery after photobleaching. PLoS One 17: e0276909

Saxton MJ (1994) Anomalous diffusion due to obstacles: a Monte Carlo study. Biophys J 66: 394–401

Schramm AC, Hocky GM, Voth GA, Blanchoin L, Martiel J-L & De La Cruz EM (2017) Actin Filament Strain Promotes Severing and Cofilin Dissociation. Biophys J 112: 2624–2633

Schramm AC, Hocky GM, Voth GA, Martiel J-L & De La Cruz EM (2019) Plastic Deformation and Fragmentation of Strained Actin Filaments. Biophys J 117: 453–463

Shin H, Purdy Drew KR, Bartles JR, Wong GCL & Grason GM (2009) Cooperativity and frustration in protein-mediated parallel actin bundles. Phys Rev Lett 103: 238102

Sinnar SA, Antoku S, Saffin J-M, Cooper JA & Halpain S (2014) Capping protein is essential for cell migration in vivo and for filopodial morphology and dynamics. Mol Biol Cell 25: 2152–2160

Suarez C, Roland J, Boujemaa-Paterski R, Kang H, McCullough BR, Reymann A-C, Guérin C, Martiel J-L, De la Cruz EM & Blanchoin L (2011) Cofilin tunes the nucleotide state of actin filaments and severs at bare and decorated segment boundaries. Curr Biol 21: 862–868

Suzuki EL, Chikireddy J, Dmitrieff S, Guichard B, Romet-Lemonne G & Jégou A (2020) Geometrical Constraints Greatly Hinder Formin mDia1 Activity. Nano Lett 20: 22–32

Svitkina TM & Borisy GG (1999) Arp2/3 complex and actin depolymerizing factor/cofilin in dendritic organization and treadmilling of actin filament array in lamellipodia. J Cell Biol 145: 1009–1026

Tamada A (2019) Chiral Neuronal Motility: The Missing Link between Molecular Chirality and Brain Asymmetry. Symmetry 11: 102

Tee YH, Goh WJ, Yong X, Ong HT, Hu J, Tay IYY, Shi S, Jalal S, Barnett SFH, Kanchanawong P, et al (2023) Actin polymerisation and crosslinking drive left-right asymmetry in single cell and cell collectives. Nat Commun 14: 776

Valencia FR, Sandoval E, Du J, Iu E, Liu J & Plotnikov SV (2021) Force-dependent activation of actin elongation factor mDia1 protects the cytoskeleton from mechanical damage and promotes stress fiber repair. Dev Cell 56: 3288–3302.e5

Vignjevic D, Kojima S-I, Aratyn Y, Danciu O, Svitkina T & Borisy GG (2006) Role of fascin in filopodial protrusion. J Cell Biol 174: 863–875

Winkelman JD, Suarez C, Hocky GM, Harker AJ, Morganthaler AN, Christensen JR, Voth GA, Bartles JR & Kovar DR (2016) Fascin- and α-Actinin-Bundled Networks Contain Intrinsic Structural Features that Drive Protein Sorting. Curr Biol

Wioland H, Ghasemi F, Chikireddy J, Romet-Lemonne G & Jégou A (2022) Using microfluidics and fluorescence microscopy to study the assembly dynamics of single actin filaments and bundles. J Vis Exp

Wioland H, Guichard B, Senju Y, Myram S, Lappalainen P, Jégou A & Romet-Lemonne G (2017) ADF/Cofilin Accelerates Actin Dynamics by Severing Filaments and Promoting Their Depolymerization at Both Ends. Curr Biol 27: 1956–1967.e7

Wioland H, Jegou A & Romet-Lemonne G (2019a) Quantitative Variations with pH of Actin Depolymerizing Factor/Cofilin’s Multiple Actions on Actin Filaments. Biochemistry 58: 40–47

Wioland H, Jegou A & Romet-Lemonne G (2019b) Torsional stress generated by ADF/cofilin on cross-linked actin filaments boosts their severing. Proc Natl Acad Sci U S A 116: 2595–2602

