## Supplementary Informations for "Fascin-induced bundling protects actin filaments from disassembly by cofilin"

**Supp. Text 1**

Impact of the detection of cofilin severing events occurring on filament bundles

The departure of one cofilin-saturated actin segment, detectable as a drop in cofilin fluorescence occurs once it has severed at its two boundaries. We expect, in the vast majority of cases, that severing occurs first at the pointed end side of cofilin clusters (Suarez *et al*, 2011; Wioland *et al*, 2017). Because of our experimental configuration, the second severing event may then occur immediately after: due to the flow, the dangling cofilin-decorated region may pivot around its barbed-end side where it is still bound by fascin to the second filament (Supp. Fig. 8). This would result in a sharp bend which would accelerate the severing at this boundary, as previously reported (Wioland *et al*, 2019b). In that case, the severing of the first cluster boundary would lead to an almost immediate detachment of the cofilin cluster from the bundle. We performed a specific assay, described in Supp. Fig. 8, allowing us to quantify that around half of the severing events occurring at the pointed end side of a cofilin cluster coincided with a drop in cofilin fluorescence (i.e. rapid departure of cofilin-actin segments).

These cofilin-actin segments will either sever at the opposite boundary, before a cofilin cluster is nucleated on the adjacent filament in the region facing the first cofilin cluster, and this will be counted as route 1 events, or a cofilin cluster will be nucleated on the adjacent filament. In the latter case, the rate of nucleation of the second cofilin cluster is the rate observed on single filaments, as there is no inter-filament cooperativity (i.e. the first cofilin cluster does not transmit any twist as it has severed at one boundary). In the competition between those two types of events, severing at the other boundary will dominate and occur ~90% of the time, considering the measured rates and a cluster size of typically ~ 100 subunits. Experimentally, these events will appear as route 1 events. The remaining 10% will ‘appear’ as route 2 events as two cofilin clusters are overlapping and as the severing of these two clusters will lead to bundle fragmentation, although they are not route 2 events per se. The error in distinguishing between route 1 and route 2 events is thus reasonably small (~10%) and does not appreciably impact the estimated rates.

Overall, the drop in cofilin fluorescence is thus a reasonable surrogate of the cofilin severing event.

**Supplementary Table 1.**

| reaction rates | values measured at 200 nM cofilin | relative to single filament rates |
|---|---|---|
| cluster nucleation on single filaments ($k_{nuc,SF}$) | $5.85.10^{-6}$ $sub^{-1}.s^{-1}$ | - |
| 1st cluster nucleation on 2-filament bundles ($k_{nuc,1}$) | $1.08.10^{-6}$ $sub^{-1}.s^{-1}$ | 1/6 |
| 2nd cluster nucleation on 2-filament bundles ($k_{nuc,2}$) | $4.7.10^{-5}$ $sub^{-1}.s^{-1}$ | 8 |
| cluster growth on 2-filament bundles ($v_{growth}$) | 1.5 $sub^{-1}.s^{-1}$ | 2/3 |
| cluster severing ($k_{sev,SF}$, $k_{sev,1}$, $k_{sev,2}$, $k_{sev,final}$) | $2.10^{-3}$ $s^{-1}$ | ~1 |

**Supplementary Fig. 1:** Cofilin binds less efficiently to larger filament bundles.

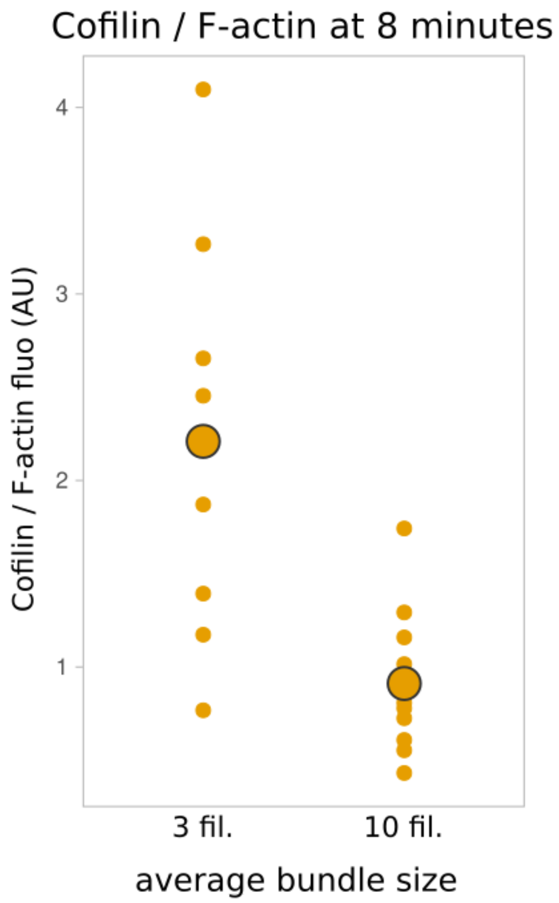


Fluorescence intensity of cofilin, normalized by the amount of F-actin, bound after 8 minutes after cofilin introduction in the ‘open chamber’, on segments of typically 4 µm of fascin-induced filament bundles of average size 3 (± 0.8, standard deviation, n = 8 segments) and 10 (± 4, n= 10 segments) filaments. The large dots represent the median value of each population.

**Supplementary Fig. 2:** Fascin crosslinking does not appear to slow down Pi release in actin filaments.

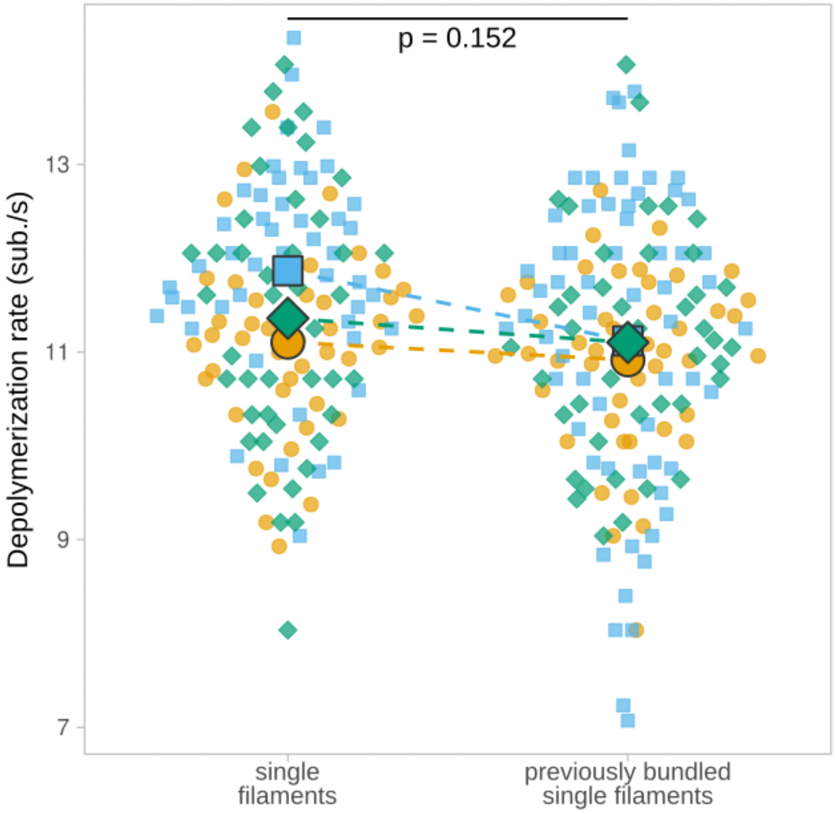


Actin filaments elongated from spectrin-actin seeds in microfluidics chambers, were bundled and aged for 15 minutes by exposing them to 0.15 µM actin and 200 nM fascin. Actin filaments are then unbundled and they depolymerize as single filaments upon exposure to buffer only. In the absence of fascin in solution, bundled filaments become individual isolated filaments after typically 30 seconds. The depolymerization rates (measured over 3 minutes) of individual actin filaments, that were initially either isolated filaments or part of 2-filament bundles when exposed to fascin, were quantified (N=3 repeats, with n = 44, 47, 44 for single filaments, and n= 45, 63, 42 for 2-filament bundles, respectively). Large symbols represent median values. The p-value is calculated from the comparison of the paired median values.

**Supplementary Fig. 3:** Fascin decreases cofilin cluster nucleation rate on single actin filaments, in a concentration dependent manner.

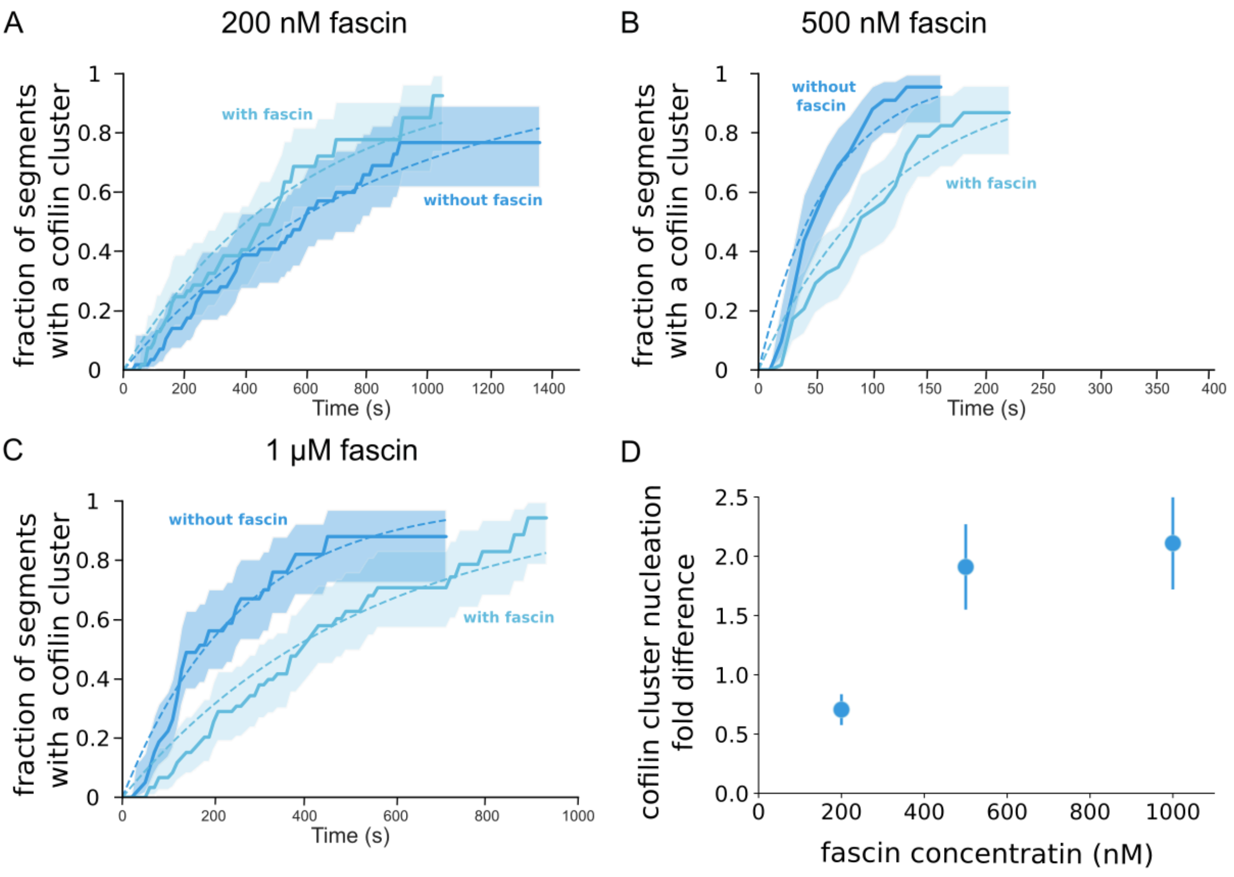


Two side-by-side populations of single actin filaments are exposed in a microfluidics chamber to 200 nM cofilin, 0.15 μM actin and either no or (A) 200 nM, (B) 500 nM, (C) 1 μM fascin (n= 54, 58, 60 segments with fascin, and n= 57, 53, 60 segments without fascin). Fraction of 5-μm segments of single actin filaments with at least one cofilin cluster, as a function of time. 95% confidence intervals are shown as shaded surfaces. Curves are fitted with a single exponential function to derive the cofilin cluster nucleation rate for each population. (D) Cofilin cluster nucleation rate fold difference between population of single actin filaments exposed to fascin or not. Error bars for each condition are 95% confidence intervals, based on the sample sizes of the two survival fractions.

.

**Supplementary Fig. 4:** Cofilin clusters nucleate homogeneously along 2-filament bundles.

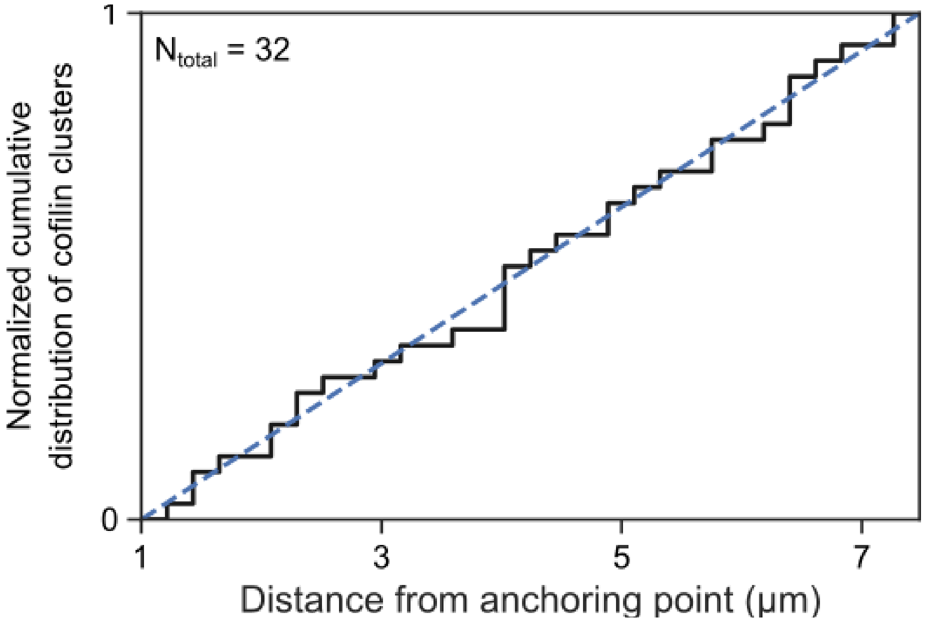


Cumulative distribution of the localization of n = 32 cofilin clusters along 7-µm 2-filament bundles, when exposed to 200 nM mCherry-cofilin-1, 200 nM fascin and 0.15 µM actin. The first micron of the bundle was excluded to avoid any effect due to curvature close to the anchorage point of the filaments. The dashed line represents a population of clusters that would be perfectly homogeneously distributed.

**Supplementary Fig. 5:** Cofilin cluster nucleation rate decreases with bundle size.

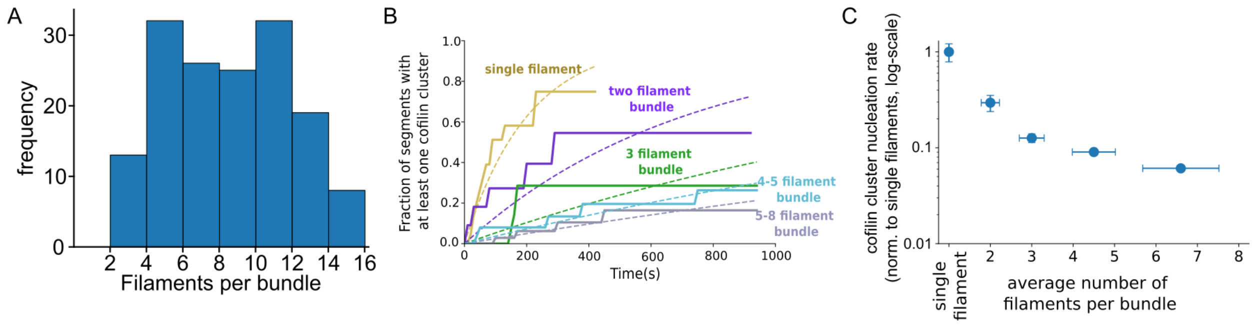


**A** Size distribution of fascin-induced bundles formed from actin filaments polymerized from individual beads in the presence of 200 nM fascin. Average size = 9.7 (± 4.8) filaments per bundle, n = 179 bundles. Size is determined by the actin fluorescent intensity relative to the intensity of single actin filaments (n = 10 filaments).

**B** Fraction of 5-µm segment filament bundles harboring at least one cofilin cluster, over time, as a function of bundle size (single filaments n = 22, 2-fil. n = 10, 3-fil. n = 20, 4-5-fil. n = 40 and 5-8-fil. bundles n = 25). Bundle sizes were determined based on the relative fluorescence intensity of actin compared to single filaments.

**C** Impact of bundle size on the cofilin cluster nucleation rate, normalized by the rate on single filaments and by the number of filaments in bundles. Values are obtained from exponential fits of curves shown in panel (A). Error bars are standard deviations.

**Supplementary Fig. 6:** Impact of fascin on the growth rates of cofilin clusters.

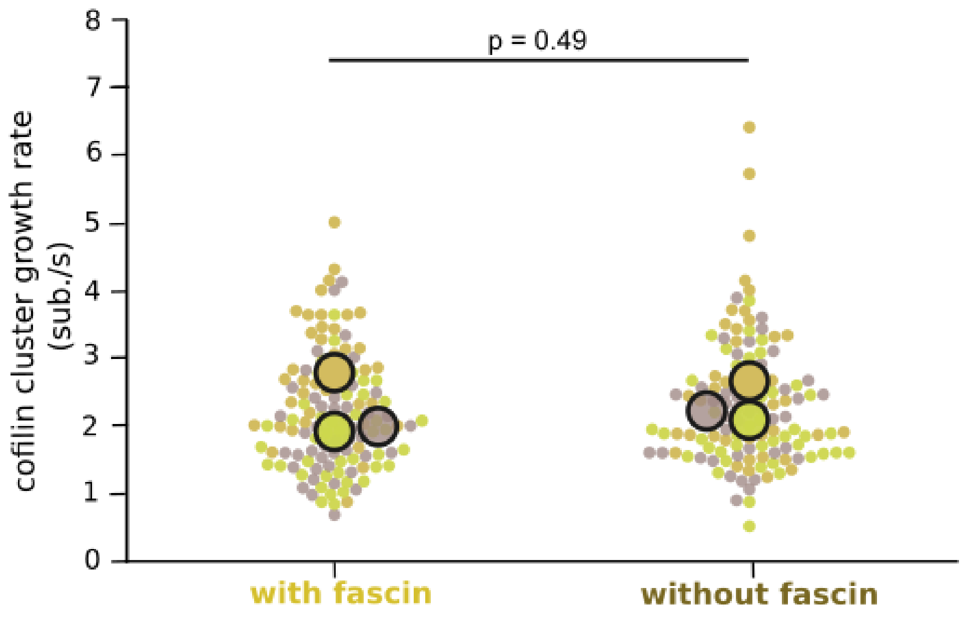


Growth rates of cofilin clusters on single filaments exposed to 200 nM mCherry-cofilin1 in the presence or absence of 200 nM fascin, N = 3 experiments, n > 40 cofilin clusters for each experiment. Large symbols represent averages over individual measures from independent experiments. Paired t-test p-value = 0.494.

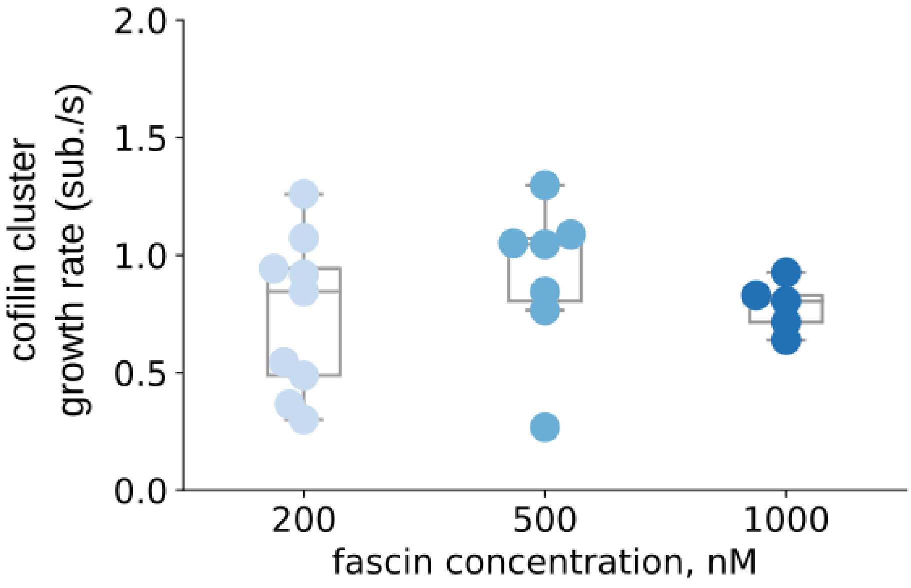


Growth rates of cofilin clusters on 2-filament bundles exposed to 200 nM cofilin and various fascin concentrations (n = 9, 7, 5 cofilin clusters for 200, 500 and 1000 nM fascin respectively). One-way ANOVA test p-value = 0.56.

**Supplementary Fig. 7:** Cofilin fluorescence reveals the presence of overlapping cofilin clusters along 2-filament bundles.

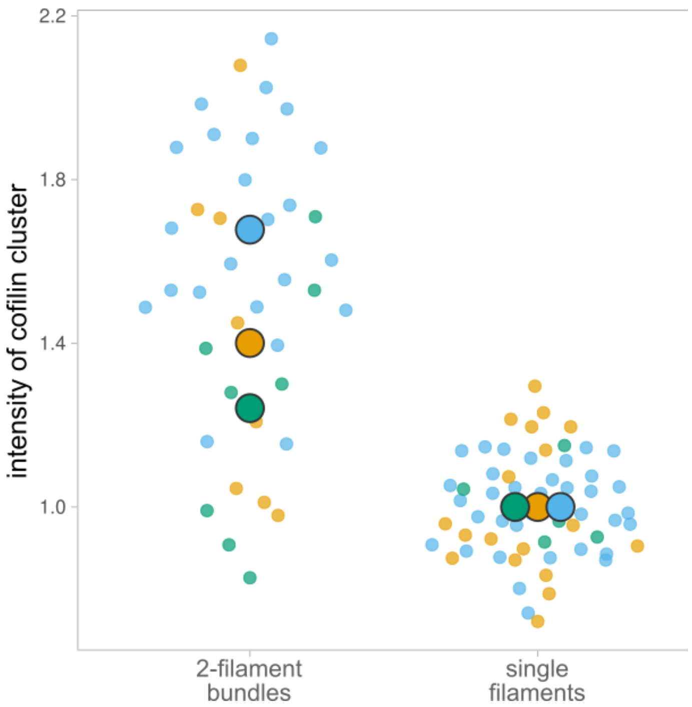


Maximum fluorescence intensity of cofilin measured over 1 μm stretches on 2-filament bundles, compared to the intensity of cofilin clusters on single actin filaments (N = 3 experiments, with n = 23 (blue), 8 (green), 8 (orange) spots analyzed on 2-filament bundles, and n = 35, 5, 18 cofilin clusters on single filaments, respectively). Large symbols represent averages over individual measures from independent experiments.

**Supplementary Fig. 8:** Curvature of cofilin-actin segments induced by the flow may accelerate severing.

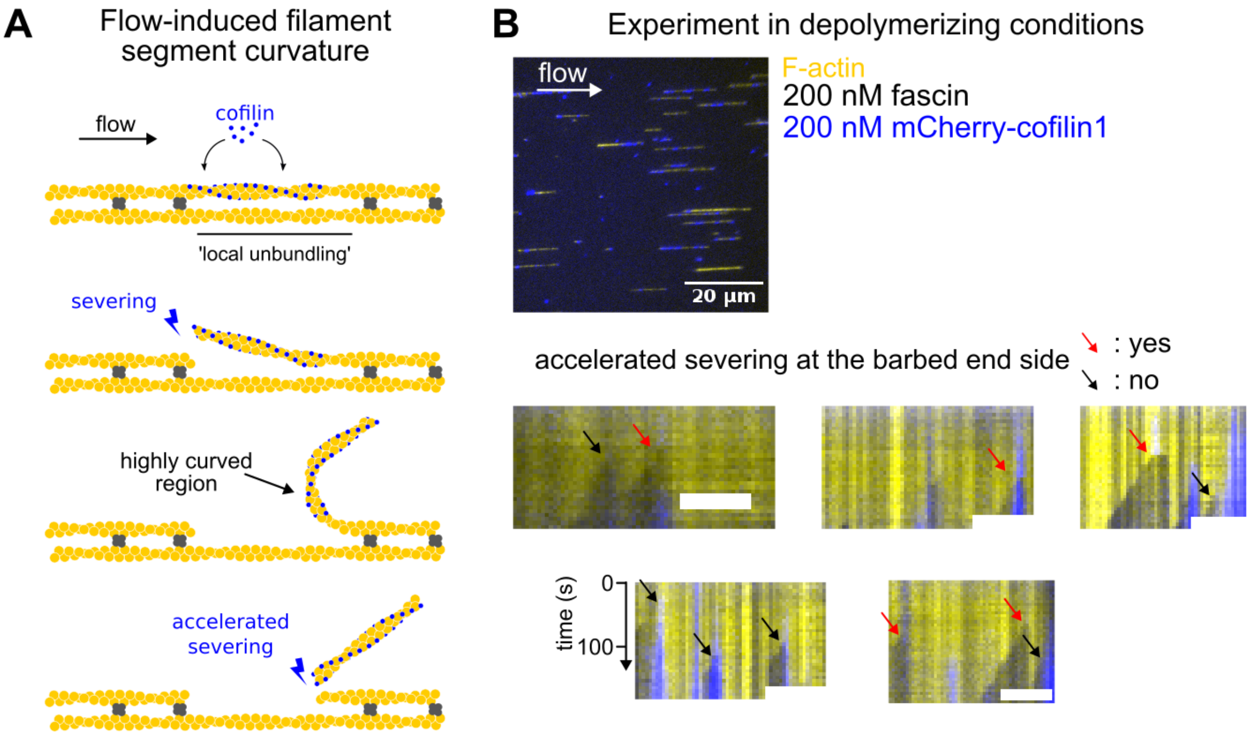


(A) Schematics illustrating how a microfluidics flow may accelerate severing at the barbed end side of a cofilin-decorated segment. Once a cofilin cluster has severed at its pointed end side, the flow may induce a high curvature at its barbed end side, which results in an increase of the severing at this boundary (Wioland *et al*, 2019b).

(B) Microfluidics assay where single filaments and 2-filament bundles are exposed to fascin and cofilin in the absence of actin. Severing at the pointed end side of cofilin clusters creates free barbed ends which depolymerize. Representative kymographs are shown (scale bar = 5 µm). Red arrows indicate pointed end side severing events which are accompanied with the departure of the cofilin cluster (i.e. due to accelerated severing at the barbed end side, as depicted in panel A), and black arrows severing events where it is not. 46% of the severing events leads to accelerated cofilin-segment departure (n = 26 events).

**Supplementary Fig. 9:** Fraction of cofilin clusters that will sever before a cofilin cluster is nucleated on a adjacent filament of a 2-filament bundle in the region facing the first cofilin cluster.

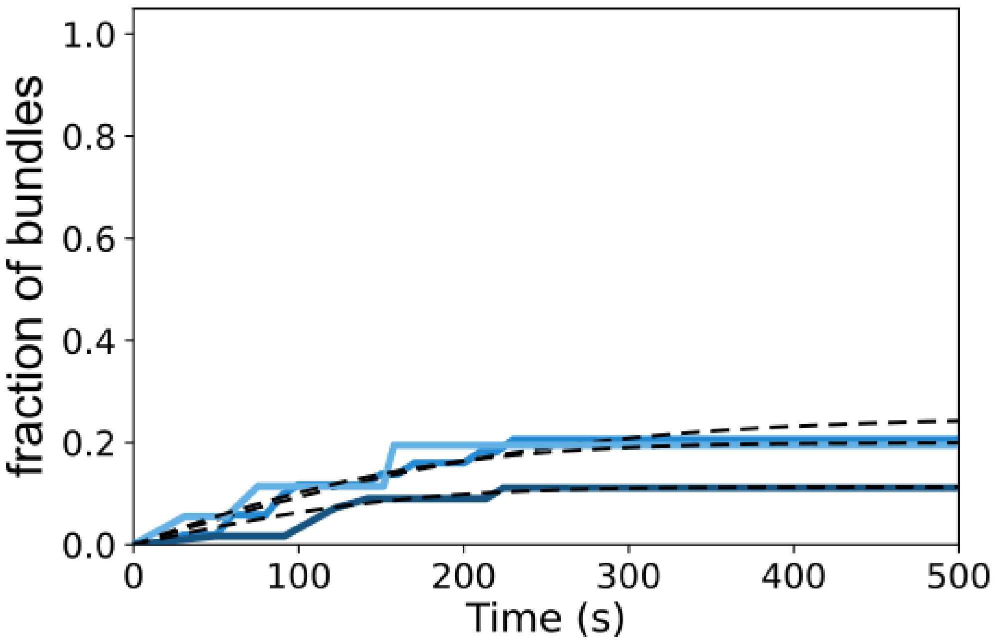


Fraction of cofilin clusters that will sever before a cofilin cluster is nucleated on a adjacent filament of a 2-filament bundle in the region facing the first cofilin cluster, as a function of time, from 3 independent experiments (n= 101,17, and 28 cofilin clusters, from top to bottom curves). Fits of the curves (dashed lines, see Methods) yield fractions of 0.3, 0.2, and 0.15.

**Supplementary Fig. 10:** Cofilin cluster severing rate on single actin filaments.

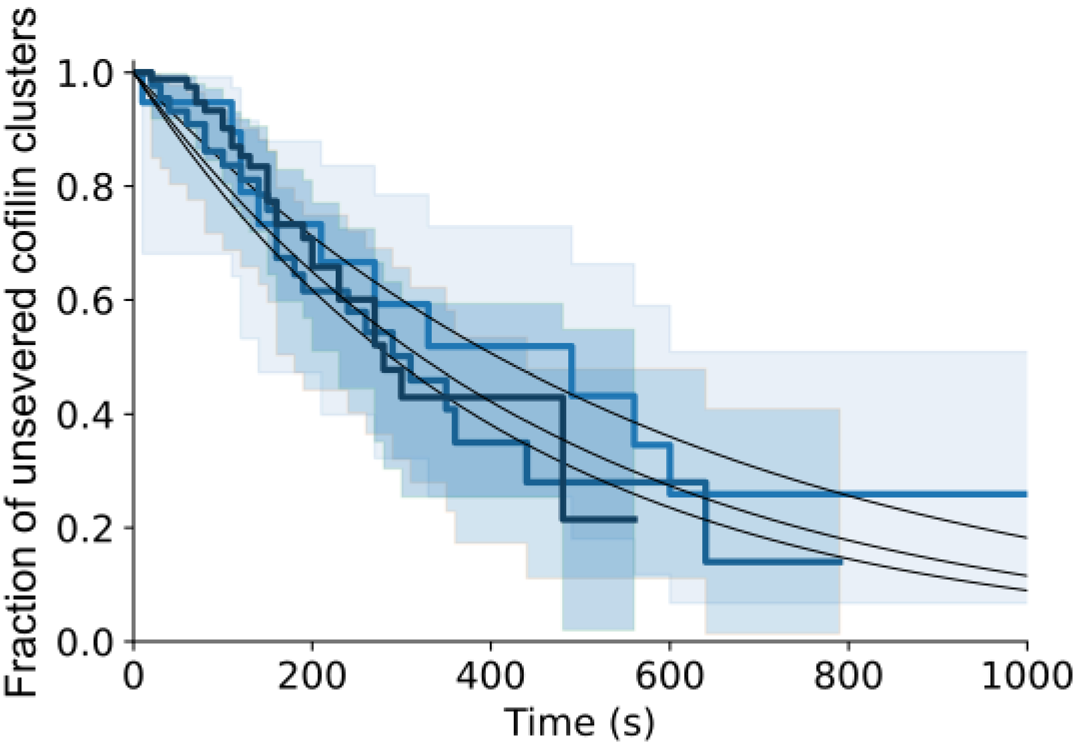


Cofilin severing rate on single actin filaments from 3 independent experiments (n= 19, 88, 45 filaments from lighter to darker blue curves). Single exponential fits (black lines) yield cofilin cluster severing rates on single filaments of 1.7, 2.15, and 2.4 .$10^{-3}$ $s^{-1}$.

**Supplementary Fig. 11:** Observations of co-localized cofilin clusters on bundles composed of more than 2 filaments.

Example 1:

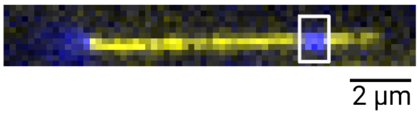


Bundle size = 3-filaments

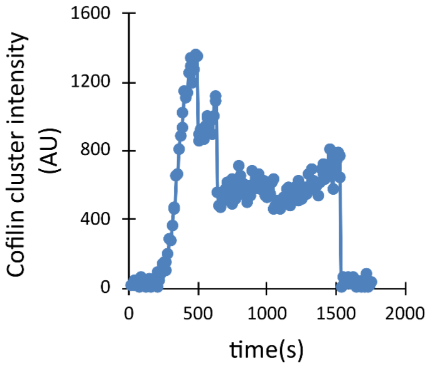


Example 2:

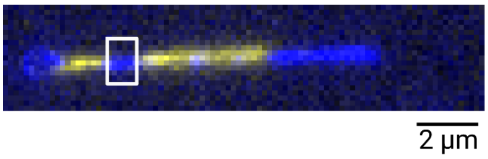


Bundle size = 8-filaments

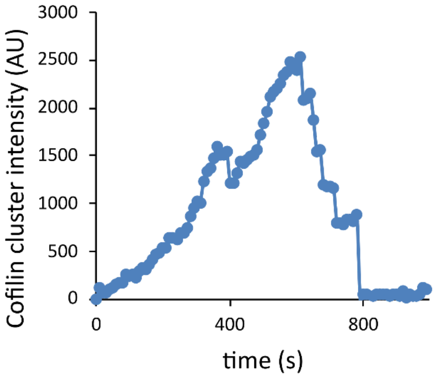


Example 3:

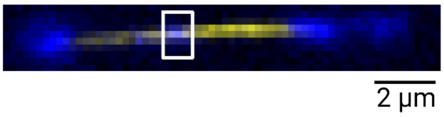


Bundle size = 5-filaments

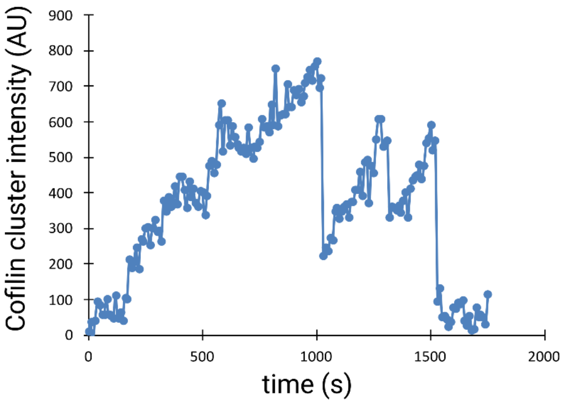


Cofilin fluorescence intensity of cofilin clusters on bundles larger than 2 filaments, showing several co-localized cofilin clusters, with multiple decreasing steps indicating cluster severing.

**Supplementary Fig. 12:** Numerical simulations indicating the impact of inter-filament cooperativity of the nucleation of cofilin clusters on the fragmentation of 2-filament bundles.

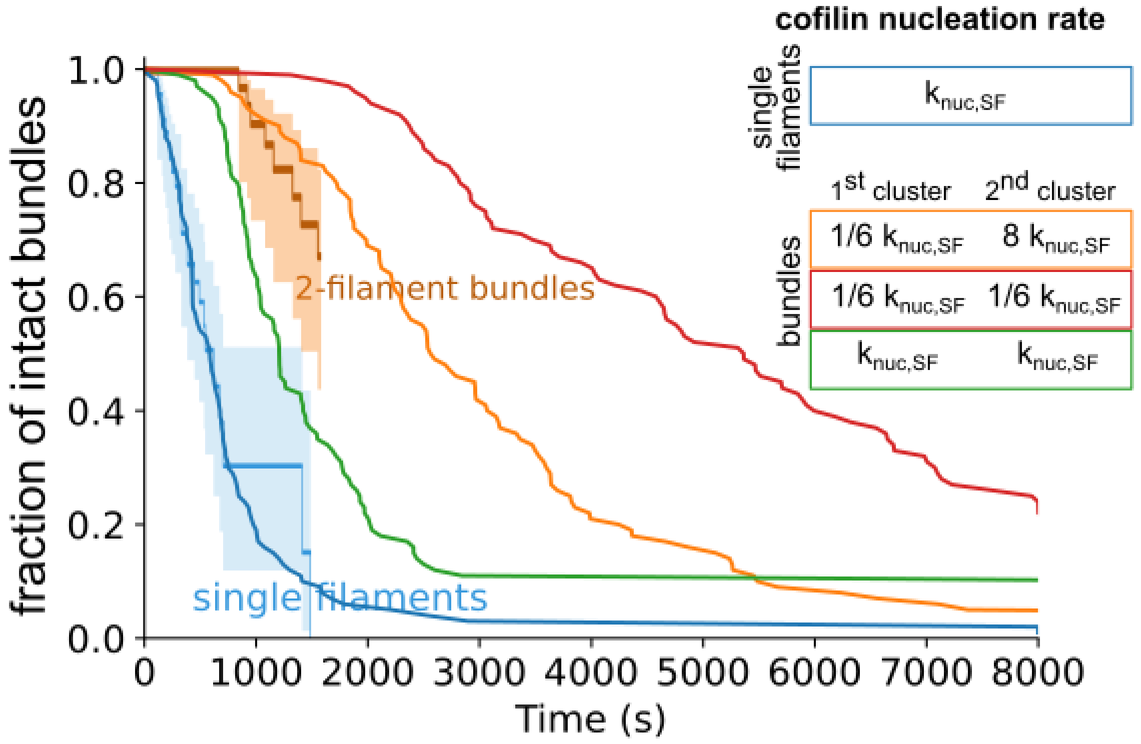


Fraction of intact single filaments or 2-filament bundles over time (n = 100 simulated 2-filament bundles for each condition) upon exposure to cofilin. The reference nucleation rate of cofilin clusters is the one on single filaments (blue curve, $k_{nuc,SF}$). On bundles, the nucleation rate of the first cofilin cluster is either the one observed experimentally for fascin-induced 2-filament bundles (orange and red curves, 1/6 $k_{nuc,SF}$) or similar to the one measured on single actin filaments (green curve, $k_{nuc,SF}$). The nucleation rate of a second cofilin cluster, i.e. on a filament in the region facing a first cofilin cluster on the other filament, is either resulting from inter-filament cooperativity imposed by fascin bundling (see Main text) (orange curve, 8 x $k_{nuc,SF}$), or not (red curve, 1/6 $k_{nuc,SF}$, and green curve, $k_{nuc,SF}$). Experimental data for single actin filaments and 2-filament bundles (from figure 2B) are shown for comparison

**Supplementary Fig. 13:** Twist-constraining 2-filament bundles leads to its faster fragmentation by cofilin.

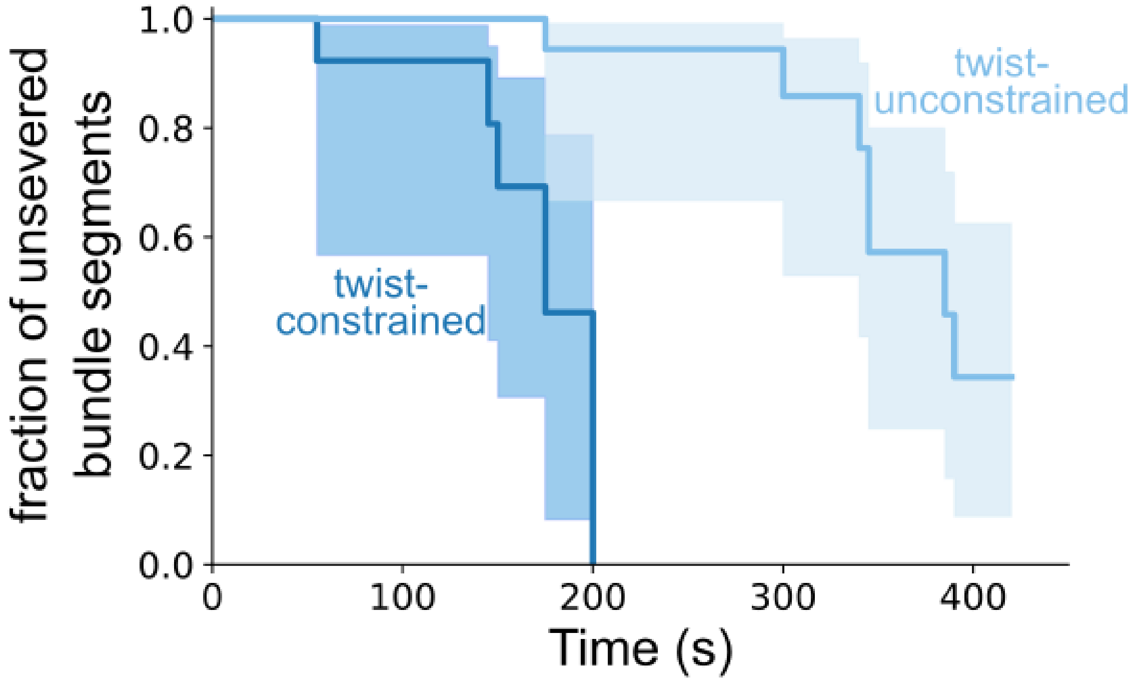


Survival fractions of unsevered 5-µm long segments from unanchored bundles (twist-unconstrained, n = 20) or anchored bundles (twist-constrained, n = 16) as a function of time, when exposed to 200 nM mCherry-cofilin1 and 200 nM fascin. 95% confidence intervals are shown as shaded surfaces. Log-rank test p-value < 0.005.

**Supplementary Fig. 14:** Cofilin cluster nucleation on twist-constrained filaments.

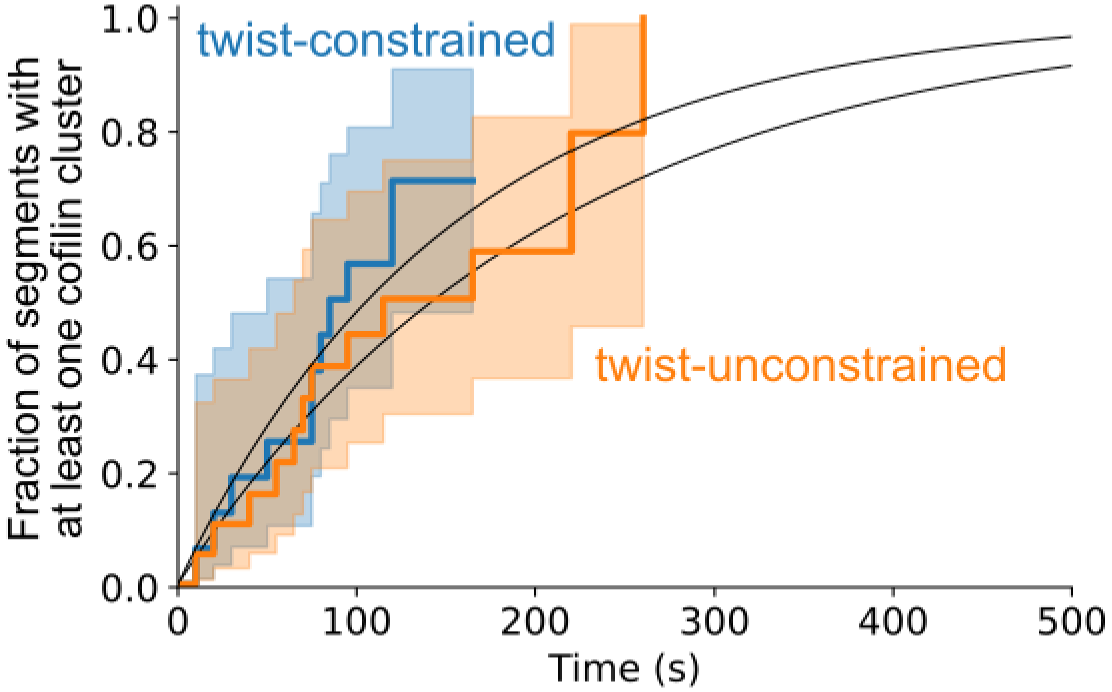


Fraction of 5-µm segments with at least one cofilin cluster on twist-constrained (n = 16 segments, blue) or twist-unconstrained (n = 20 segments, orange) bundles. Log-rank test p-value = 0.45.

**Supplementary Fig. 15:** Numerical simulations showing the impact of inter-filament cooperativity cofilin cluster nucleation on the fragmentation of twist-constrained 2-filament bundles.

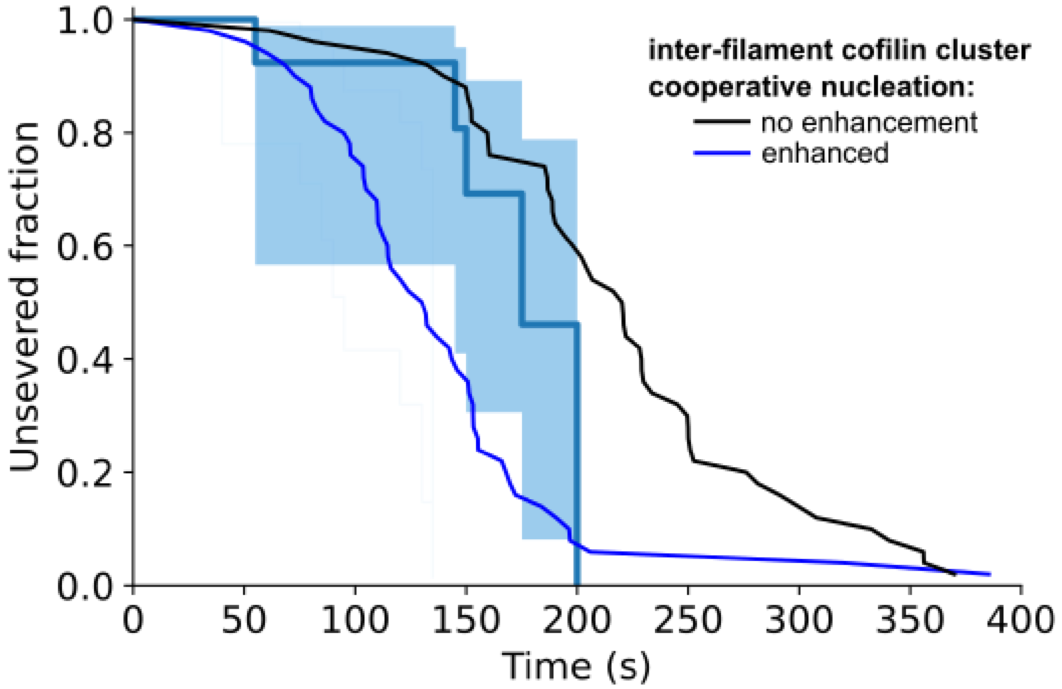


Results of numerical simulations showing that fragmentation of twist-constrained 2-filament bundles (n = 50 simulated 2-filament bundles) is faster with inter-filament cofilin cluster cooperative nucleation than without. Simulations with no inter-filament cooperativity seem to better reflect the experimental observations (data from Fig. 7C, n = 16 segments of 2-filament bundles).
